# Dopamine D2 Receptors modulate hippocampal theta rhythm through somatostatin interneurons

**DOI:** 10.1101/2023.05.22.541716

**Authors:** Pola Tuduri, Emmanuel Valjent, Jeanne Ster

**Author notes:** Contributed equally (Emmanuel Valjent, Jeanne Ster). Correspondence should be addressed to Pola Tuduri and Jeanne Ster.

## Abstract

Spatiomolecular mapping of hippocampal dopamine receptor D2 (D2R) neurons revealed that in addition to hilar mossy cells, hippocampal SST-INs and, to a lesser extent, PV-INs expressed D2R. However, while the role of D2R signaling onto mossy cells has been characterized, the consequence of D2R activation onto hippocampal SST-INs and PV-INs is unknown. By combining pharmacological approaches and patch-clamp recordings in organotypic hippocampal slices from control or mice lacking *Drd2* selectively in SST-INs and PV-INs, we found that D2R activation increase excitability in SST-INs while it decreases in PV-INs. These D2R-mediated changes rely on distinct intracellular pathways, involving the non-canonical β- arrestin-dependent pathway in SST-INs and the G protein-dependent pathway in PV-INs. Finally, our study also unveiled that D2R activation modulates theta oscillations through SST- INs D2R signaling.

## Introduction

The hippocampus formation is involved in a wide range of functions including motivation, learning and memory. This structure is finely tuned by numerous neuromodulator systems notably monoamines. Among these, dopamine (DA) stands out as crucial role in the cognition and hippocampal information processing ^1–3^. It is well known that DA contributes in a significant way to spatial learning ^4, 5^ as well as detection of novel and salient stimuli ^6, 7^. Disruption of hippocampal dopaminergic function has been associated with core symptoms of several neurological and neuropsychiatric including schizophrenia ^8, 9^, depression ^10^, Huntington’s diseases ^11^, Parkinson ^12^, attention-deficit hyperactivity disorders ^13–15^, and Alzheimer’s diseases ^16–18^.

A growing body of evidence indicate the role of dopamine receptor 2 (D2R) regulates basal synaptic transmission and synaptic plasticity, which are believed to be at the root of information processing and mnesic functions. A wealth of investigations show that in CA1 region, pharmacological blockade or genetic invalidation of D2R lead to a reduction in LTP and its maintenance ^3, 19–25^. Moreover, D2R activation can also impact the excitability of the CA1 network ^26, 27^. Nevertheless, the interpretation of these results is questionable given the absence of D2R in pyramidal cells and reflect a lack of knowledge about the localization, molecular identity, intracellular signaling pathways, and function of D2R-expressing cells in the hippocampus.

The development of transgenic mouse lines combined with immunohistochemistry or *in situ* hybridization have revealed the specific cellular localization of D2R-containing cells ^28–30^. Discrete sub-populations of GABAergic cells expressing D2R were detected in the rostral hippocampus in the CA3/CA1 areas ^29^. D2Rs were mainly expressed in two different subtypes of GABAergic cells: somatostatin (SST) and parvalbumin (PV) cells that can finely regulate the hippocampus network by innervating specific and distinct subcellular domains of pyramidal cells ^31^. While our knowledge of the identity of D2R receptor-expressing neurons is being refined, the role of these receptors in these different sub-types of interneurons remains poorly understood.

In this context, to clarify the role of hippocampal dopaminergic transmission via the recruitment of D2Rs in these two classes of interneurons, we performed whole-cell patch-clamp recordings in organotypic hippocampal slice model. We assessed the effect of D2R activation on local neuronal excitability as well as hippocampal oscillatory activity by selectively suppressing D2R in SST+ or PV+. Our results unveiled a key role of SST-INs D2R/β-arresting signalling in the modulation of theta oscillations.

## Materials and Methods

### Animals

*tdTomato*^SST^ and *tdTomato*^PV^ mouse lines were generated by breeding Rosa26-*tdTomato* mice (Jax Stock# 007909) with Sst-IRES-cre mice (JaxStock#013044) or Pvalb-cre mice (B6;129P2- Pvalbtm1(cre)Arbr/J). *Drd2*^SST^ and *Drd2*^PV^ mouse lines were generated by breeding *Drd2^loxP/loxP^* (B6.129S4(FVB)-Drd2tm1.1Mrub/J) with *Sst-IRES-Cre* or *Pvalb-Cre* mice respectively. Male C57Bl/6 mice were purchased from Charles River Laboratories (France). Animals were housed under standardized conditions with a 12 h light/dark cycle, ad libitum food and water, stable temperature (22 ± 2°C) and controlled humidity (55 ± 10%). Housing and experimental procedures were approved by the French Agriculture and Forestry Ministry (A34- 172-13). The animal care and experimental procedures were performed in accordance with the animal welfare guidelines 2010/63/EC of the European Communities Council Directive.

### Organotypic hippocampal slices

Organotypic hippocampal slices were prepared from 6-day-old mice pups as previously described ^32^ following a protocol approved by the veterinary Department of Animal Care of Montpellier. Three to five mice were used for each slice culture preparations and each experimental condition was tested over the course of at least three preparations. Slices were placed on a 30 mm porous membrane (Millipore, Billerica MA, USA) and kept in 100 mm diameter Petri dishes filled with 5 ml of culture medium containing 25% heat-inactivated horse serum, 25 % HBSS, 50 % Opti-MEM, penicillin 25 units/ml, streptomycin 25 μg/ml (Life technologies). Cultures were maintained in a humidified incubator at 37°C and 5 % CO_2_ until DIV3 (day *in vitro*) and then were kept at 33 °C and 5 % CO_2_ until the electrophysiological experiments.

### Electrophysiology

After 3 weeks *in vitro*, slices cultures were transferred to a recording chamber on an upright microscope (Olympus, France). Slices were superfused continuously at 1 ml/min with a solution containing the following (in mM) 125 NaCl, 2.7 KCl, 11.6 NaHCO_3_, 0.4 NaH_2_PO4, 1 MgCl_2_, 2 CaCl_2_, 5.6 D-glucose, and 0.001% phenol red (pH 7.4, osmolarity 305 mOsm) at 31°C. Whole cell recordings were obtained from cells held at −70 mV using a Multiclamp 700B amplifier (Axon Instruments, Union City, CA, USA). Recording electrodes made of borosilicate glass had a resistance of 4-6 MΩ (Warner Instruments, USA) and were filled with K-gluconate solution containing (in mM) 125 K-gluconate, 5 KCl, 10 Hepes, 1 EGTA, 5 Na-phosphocreatine, 0.07 CaCl_2_, 2 Mg-ATP, 0.4 Na-GTP (pH 7.2, osmolarity 300 mOsm). Membrane potentials were corrected for junction potentials. For all recording conditions, only cells with access resistance < 20 MΩ and a change of resistance < 25% over the course of the experiment were analyzed. Data were filtered with a Hum Bug (Quest Scientific, Canada), digitized at 2 kHz (Digidata 1444A, Molecular Devices, Sunnyvale, CA, USA), and acquired using Clampex 10 software (Molecular Devices).

To assess intrinsic excitability of the neuron, increasing depolarizing current pulses (ranging from 0 to + 360 pA in 60 pA increments, 3000 ms duration) were injected into the neuron. The number of action potentials (AP) was calculated for the second step after the initiation of the first action AP. For the latency to spike and the interval inter spike, we used the first step with AP. quinpirole was perfused at 1 µM and 10 µM. For experiments using C57Bl/6J mice, somatostatin and parvalbumin cells were patched in the *stratum oriens*. They can be separated by electrophysiological response notably during injection of hyperpolarizing current (−200 pA for 3000 ms) and the firing rate. To confirm the identity of the cell, biocytin was added into the recording pipette for post hoc analysis of morphological features.

For the signaling pathway experiment, a GSK3 inhibitor was added to the intracellular medium (SB216763, 1 mM) while pertussis toxin (PTX, 500 nM) was incubated in the culture medium at least 18h before starting the experiment.

Theta oscillations were filtered at 2–5 kHz and analyzed off-line (pCLAMP 10; Axon Instruments,^32^). Oscillation analyses were performed after 20 min of application of MCh with or without quinpirole (10 µM) or raclopride (40 µM) during one minute. Calculation of oscillatory activity was performed from the time of the second peak in the Clampfit autocorrelation function. A segment was considered rhythmic when the second peak of the autocorrelation function was at least 0.3 and several regularly spaced peaks appeared.

The Potassium currents (K^+^) were examined in somatostatin and parvalbumin neurons as previously described ^33^. Briefly, we recorded somatostatin and parvalbumin cells in the CA1 area in voltage clamp mode. BAPTA (5 mM) was included in the internal recording solution. The K^+^ currents were isolated with the addition of 0.1 µM TTX, 10 µM XE 991 dihydrochloride, 0.1 mM ZD2788, 50 µM d-AP5, 10 µM NBQX, 100 µM CGP3548, and 3 µM of gabazine in the extracellular solution. After a pre-pulse to −70 mV, we applied 300 ms voltage steps every 10 mV (−70 to −10 mV) and every 10 mV (−100 to −80 mV). Leak currents were measured from the second series, extrapolated to the first series, and subtracted. Current amplitude was measured in the last 50 ms of the 300 ms pulse. To separate Kv1.1 current from other delayed rectifier currents, dendrotoxin-K (DTX-K, 100 nM) was washed in, and DTX- insensitive currents were subtracted from the baseline current.

### Drugs

Aripiprazole, 1,2-Bis (2-Aminophenoxy) ethane-N, N, N′, N′-tetraacetic acid (BAPTA), Bovine Serum Albumin (BSA), Dimethyl sulfoxide (DMSO), Acetyl-β-methylcholine chloride (metacholine) and Pertussis toxin (PTX) were purchased from Sigma-Aldrich. CGP55845 hydrochloride (CGP55845), D-2-aamino-5-phopshonovalerate (D-AP5), SR 95531 hydrobromide (Gabazine), NBQX disodium salt (NBQX), (-)-Quinpirole hydrochloride (Quinpirole), Raclopride, Tetrodotoxin citrate (TTX), XE 991 dihydrochloride (XE 991), ZD- 7288 were purchased from Tocris. Dendrotoxin-K (DTX- K) and SB-216763 were purchased from Alomone lab and Hellobio respectively.

### Statistical analysis

GraphPad Prism v7.0 software was used for statistical analyses. Data are shown as the means ± SEM. For normally distributed parameters, Student’s *t* test (paired or unpaired, two-sided) was used to compare to sets of data. Multiple comparisons were performed by one-way or two-way ANOVA repeated measures followed by a post-hoc analysis. * p < 0.05, ** p < 0.01, *** p < 0.001 and **** p < 0.001. Values are indicated in the Supplemental Table.

## Results

### Opposing intrinsic electrophysiological responses in hippocampal CA1 somatostatin-and parvalbumin-expressing GABAergic neurons in response to D2R activation

To determine the role of D2R on CA1 somatostatin-and parvalbumin-interneurons (named thereafter SST-INs and PV-INs), patch-clamp recordings were performed using organotypic hippocampal slices from C57Bl/6J mice. We first identified SST-INs and PV-INs based on well-established electrophysiological (Firing rate, Ih current and rebound spikes following hyperpolarized step) and morphological properties ^31, 34^. Biocytin-filled recorded neurons also confirmed the morphological features of SST-INs and PV-INs (**Figure 1a, b**). We then investigated the effect of a D2R-like agonist, quinpirole, on CA1 SST-INs and PV-INs excitability assessed by measuring the firing rate of cells in response to a series of depolarizing current steps (60 pA). In SST-INs, bath application of quinpirole (1 or 10 µM, 10 min) increased the number of action potentials (**Figure 1c, d**). This effect was observed in 7 out of 10 of recorded cells (responsive SST-INs) suggesting that a fraction of SST-INs might not express D2R (non-responsive SST-INs) (**Figure 1c, d**). Conversely, bath application of quinpirole 1 µM but not 10 µM decreased the number of action potentials in responsive PV-INs (3 out of 10), which were outnumbered by PV-INs non-responsive cells (7 out of 10) (**Figure 1e, f**). To unambiguously identify SST-INs and PV-INs in hippocampal CA1 area, similar patch-clamp recordings analyses were performed in organotypic hippocampal slices from *tdTomato*^SST^ and *tdTomato*^PV^ mice expressing the red fluorescent protein *tdTomato* in PV-and SST-expressing cells respectively. As previously observed, bath application of quinpirole increased SST-INs excitability (**Figure 1g, i, j**) and decreased PV-INs excitability (**Figure 1h, k, l**). Bath application of quinpirole at 10 µM also produced a transient decreased of excitability in PV-INs (**Supplemental Figure 1**). Importantly, the proportion of responsive *vs.* non-responsive SST-INs and PV-INs was similar between *tdTomato*^SST^/C57Bl/6J and *tdTomato*^PV^/C57Bl/6J mice. Altogether, these results indicate that D2R activation triggers opposite effects on intrinsic electrophysiological responses in hippocampal CA1 SST-INs and PV-INs.

**Figure 1:**
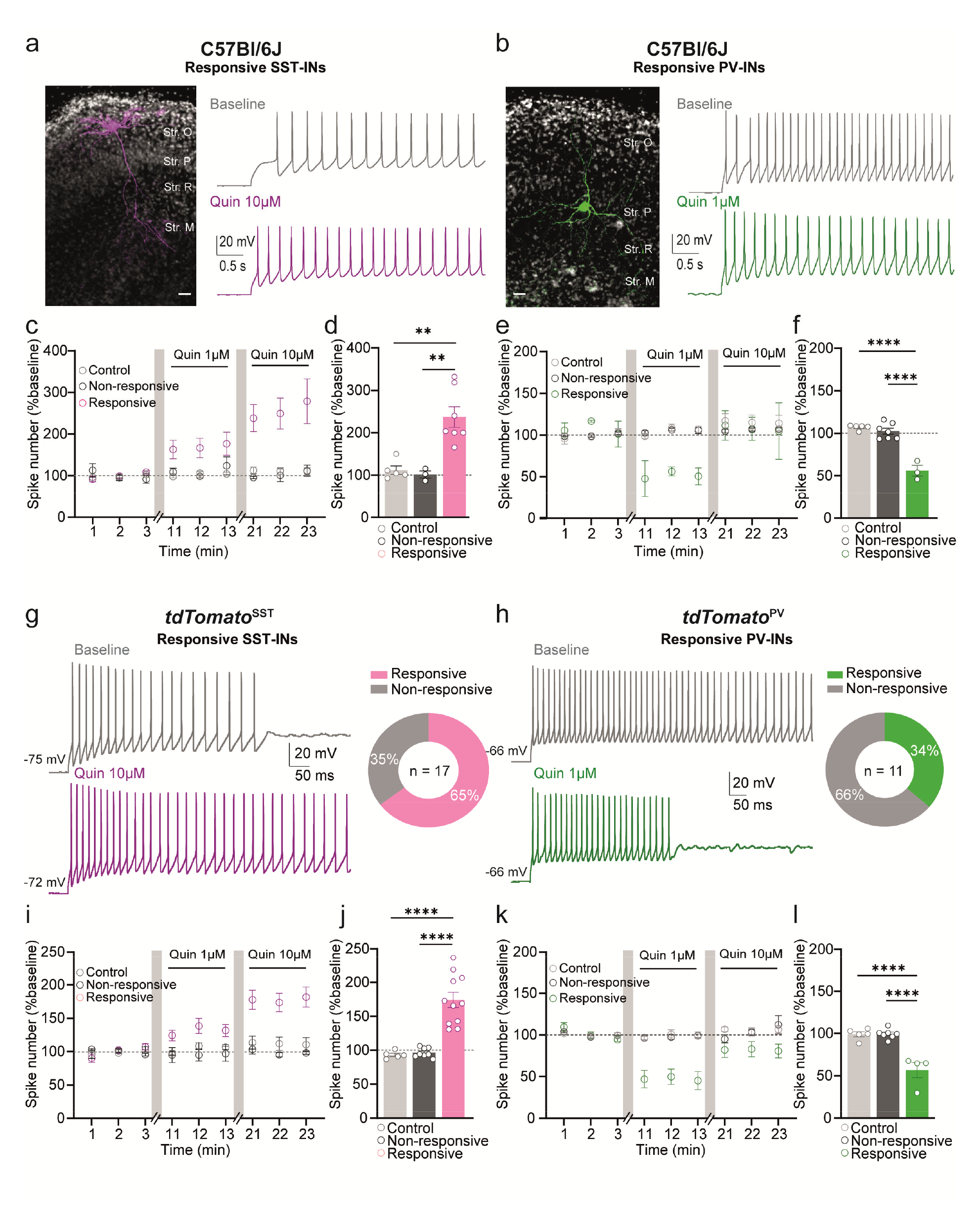
Opposing intrinsic electrophysiological responses in hippocampal CA1 somatostatin-and parvalbumin-expressing GABAergic neurons in response to D2R activation. **(a)** Upper Left, biocytin-filled responsive CA1 SST-INs from a C57Bl/6J mouse organotypic slice (scale bar: 100 µm). Upper Right, example of the firing traces of the responsive SST-INs in the presence or absence of quinpirole. (**b** as in **a**) for CA1 PV-INs from C57Bl/6J mouse. (**c**) Time course of responses to depolarizing current steps prior and in response to quinpirole 1µM and 10 µM of SST-INs. (**d**) Mean responses corresponding to the last three min epoch of quinpirole 10 µM. (**e** & **f** as in **c** & **d**) Respectively for CA1 PV-INs. (**g**) Left, example of quinpirole-responsive SST firing traces from the organotypic *tdTomato*^SST^ slice in the absence or presence of quinpirole (10 µM). Right, pie chart depicting proportions of non-responsive and responsive SST-INs to quinpirole (n = 17 cells). (**i** & **j** as in **c** & **d**) for CA1 SST-INs from *tdTomato*^SST^. Left, example of quinpirole-responsive PV firing traces from the organotypic *tdTomato*^PV^ slice in the absence or presence of quinpirole (1 µM). Right, pie chart depicting proportions of non-responsive and responsive PV-INs to quinpirole (n = 11 cells). (**k** & **l** as in **i** & **j**) for PV-INs from *tdTomato*^PV^. Data are represented as mean ± SEM. One-way ANOVA followed by Tukey test. ** p < 0.01, **** p < 0.0001; Str. M: stratum lacunosum-moleculare; Str. R: stratum radiatum; Str. P, stratum pyramidale; Str.O, stratum oriens.

### D2R activation changes intrinsic excitability of SST-INs and PV-INs through the recruitment of delayed rectifier K channels

We then analyzed whether resting membrane potential, input resistance, conductance, action potential threshold, spike afterhyperpolarisation (AHP), and action potential half-width were altered by quinpirole in responsive SST-INs and PV-INs. As detailed in **Table 1** and **2,** none of these parameters were modified indicating that alteration in the passive membrane properties may not account for changes of SST-INs and PV-INs excitability induced by quinpirole.

**Table 1.** Summary of passive properties of SST-INs in presence of quin 10µM. Data are presented as the mean ± SEM

| Parameter | Control | Responsive | Non-responsive |
| --- | --- | --- | --- |
| Resting Potential | -65,00 ± 1,924 | -63,09 ± 1,569 | -65,00 ± 1,483 |
| Rin (MΩ) | 97,82 ± 17,06 | 103,3 ± 11,78 | 105,4 ± 12,95 |
| Conductance (nS) | 3,225 ± 0,3079 | 3,317 ± 0,2531 | 3,051 ± 0,4314 |
| Capacitance (nF) | 77,82 ± 7,42 | 105,1 ± 11,35 | 103,3 ± 11,78 |
| Tau (ms) | 25,73 ± 1,492 | 29,34 ± 2,045 | 27,68 ± 3,795 |
| Peak AP (mV) | 70,40 ± 5,205 | 64,48 ± 2,571 | 68,85 ± 5,159 |
| Half AP (ms) | 1,025 ± 0,1668 | 0,8651 ± 0,08731 | 1,113 ± 0,1680 |
| AHP (mV) | -15,63 ± 4,483 | -15,47 ± 3,148 | -14,64 ± 1,624 |

**Table 2.**
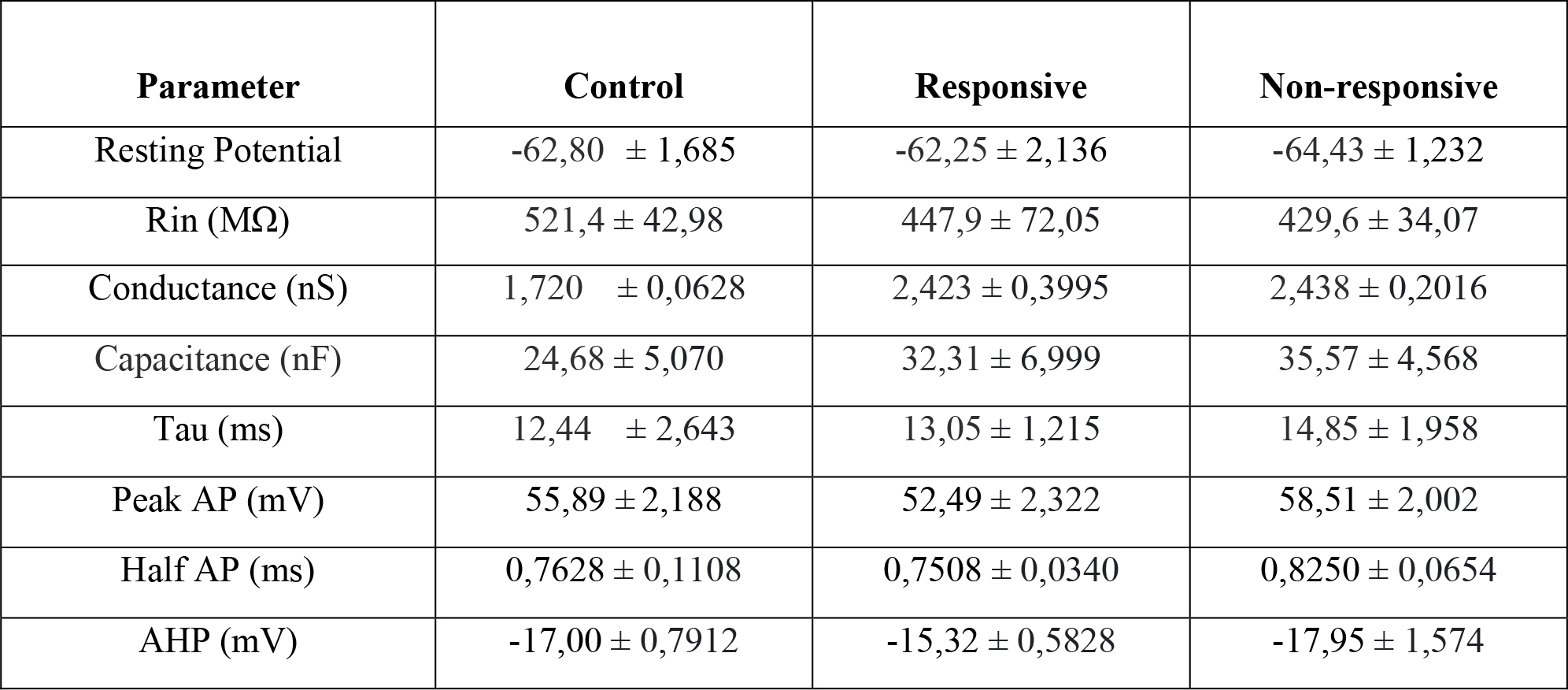
Summary of passive properties of PV-INs in presence of quin 1µM. Data are presented as the mean ± SEM

| Parameter | Control | Responsive | Non-responsive |
| --- | --- | --- | --- |
| Resting Potential | -62,80 ± 1,685 | -62,25 ± 2,136 | -64,43 ± 1,232 |
| Rin (MΩ) | 521,4 ± 42,98 | 447,9 ± 72,05 | 429,6 ± 34,07 |
| Conductance (nS) | 1,720 ± 0,0628 | 2,423 ± 0,3995 | 2,438 ± 0,2016 |
| Capacitance (nF) | 24,68 ± 5,070 | 32,31 ± 6,999 | 35,57 ± 4,568 |
| Tau (ms) | 12,44 ± 2,643 | 13,05 ± 1,215 | 14,85 ± 1,958 |
| Peak AP (mV) | 55,89 ± 2,188 | 52,49 ± 2,322 | 58,51 ± 2,002 |
| Half AP (ms) | 0,7628 ± 0,1108 | 0,7508 ± 0,0340 | 0,8250 ± 0,0654 |
| AHP (mV) | -17,00 ± 0,7912 | -15,32 ± 0,5828 | -17,95 ± 1,574 |

The regulation of intrinsic excitability relies in part on the properties and distribution of ion channel expression levels ^35^. To determine the consequence of D2R activation on spike thresholds in SST-INs and PV-INs, we examined the latency of the first spike as well as the inter-spike interval (ISI) of the first 2 action potentials (ISI being the time duration between two successive spikes in a spike train; ^36^). As show in **Figure 2a-c**, quinpirole (10 µM) decreases the first spike latencies (**Figure 2a-b** and **Supplemental Figure 2a**) as well as the ISI for the first action potential only in responsive SST-INs (**Figure 2c** and **Supplemental Figure 2a**). Opposite effects were observed in responsive PV-INs. Indeed, quinpirole (1 µM) increases the latency to spike (**Figure 2d-e** and **Supplemental Figure 2b**) and the ISI (**Figure 2f** and **Supplemental Figure 2b**).

**Figure 2:**
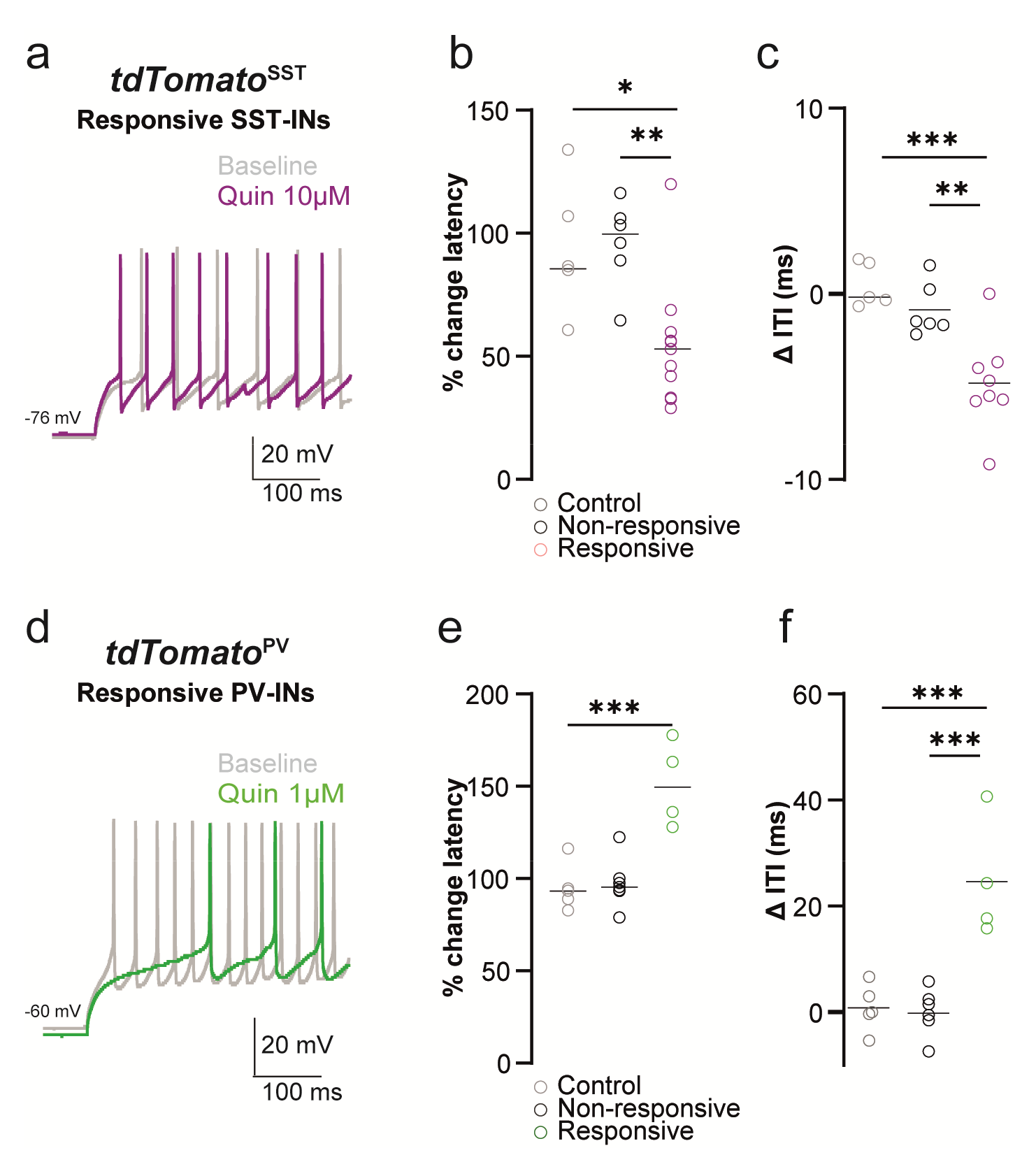
Consequence of D2R activation on active membrane properties of SST-INs and PV-Ins. (**a-c**) *tdTomato*^SST^ mouse line (quinpirole 10 µM). (**a**) Example trace showing a decrease of spike latency and ITI after quinpirole application in a responsive SST-INs. (**b**) Comparison of quinpirole-mediated latency change, (**c**) of quinpirole-mediated variability of ITI between non-responsive, responsive and control groups. (**d-f as in a-c respectively**) *tdTomato*^PV^ mouse line (quinpirole 1 µM).. Data are represented as mean ± SEM. One-way ANOVA followed by Tukey test. * p < 0.05, ** p < 0.01, *** p < 0.001.

Spike thresholds regulation, in particular the latency to spike, is intimately linked to delayed-rectifier K^+^ channels, a Kv channel subpopulation comprising the Kv1 and Kv2 family ^33, 37^. To determine whether quinpirole-induced alteration of SST-INs and PV-INs excitability relied on modulation of K^+^ conductance, we measured the delayed-rectifier K^+^ current using a whole-cell voltage-clamp protocol to isolate Kv1 and Kv2 channels based on their activation thresholds, the Kv1 family being activated from −40 mV while the Kv2 family from −30 mV^33^ (**Figure 3**). Bath application of quinpirole (10 µM) significantly decreased the K^+^ current (steady-state current) in responsive SST-INs compared to control or non-responsive SST-INs at both −40 mV and −30mV (**Figure 3b, c**). Although the leak conductance was not altered by quinpirole, we noted that responsive SST-INs were characterized by a lower leak conductance than non-responsive SST-INs (**Figure 3d**) suggesting that responsive and non-responsive SST-INs are molecularly and functionally distinct. In contrast, bath application of quinpirole (1 µM) increased K^+^ currents in responsive PV-INs at both −40 and −30 mV (**Figure 3e-g**) without altering the leak conductance (**Figure 3h**).

**Figure 3:**
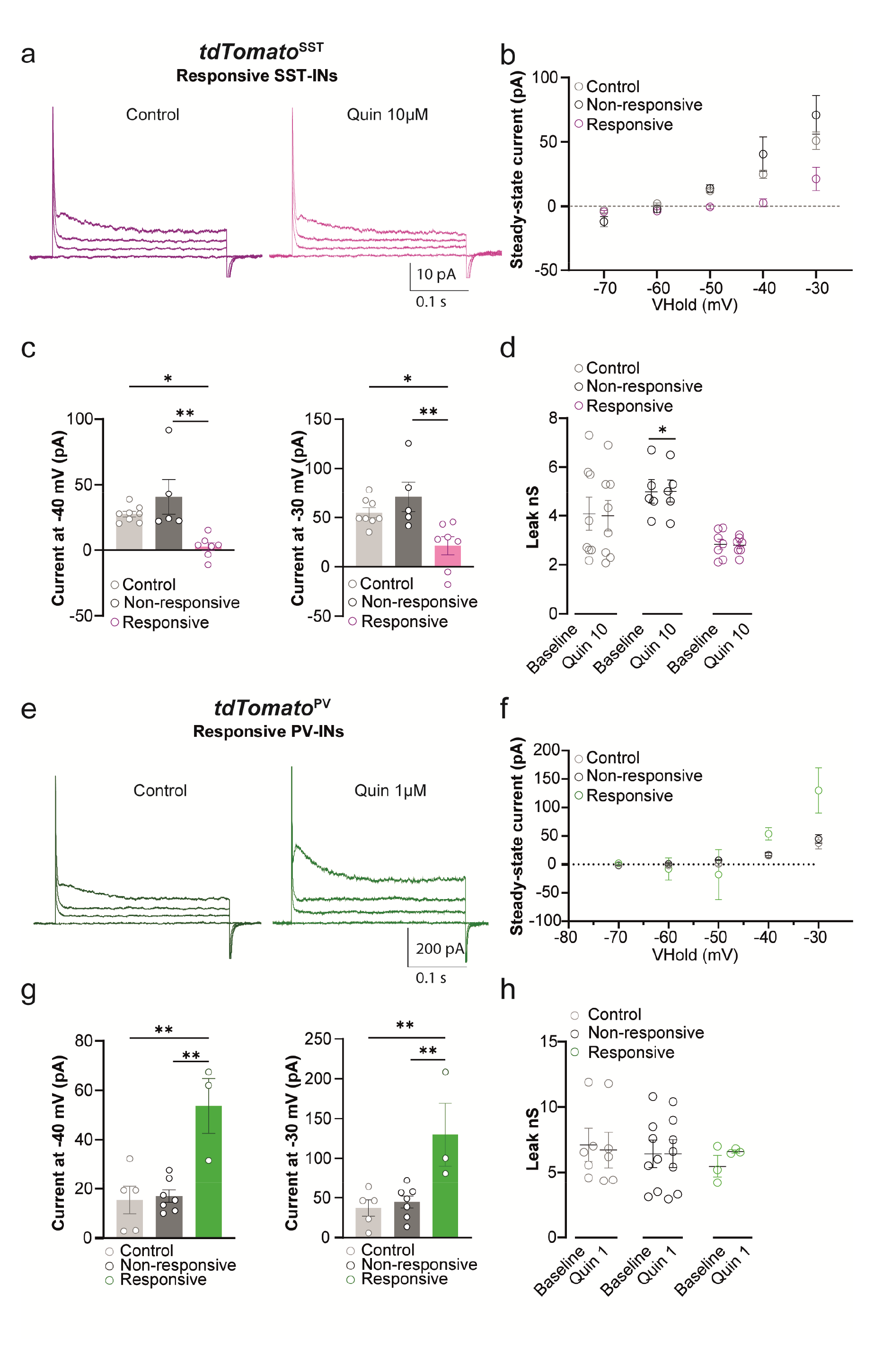
Changes in active membrane properties mediated by D2R activation are underpinned by K^+^ currents in SST-INs and PV-INS. (**a-d**) Whole-cell patch clamp recording from *tdTomato^SST^* cells. (**a**) Voltage-clamp protocol to measure delayed rectifier K currents, and example trace currents from one responsive SST-INs. (b) I–V curve of leak-subtracted current for SST-INs in organotypic slices from −70 to −30 mV. (c) Mean of leak-subtracted steady state at −40 mV or −30mV in SST-INs. (**d**) Leak conductance prior to and after quinpirole 10 µM between responsive, non-responsive and control groups. (**e-h**) Whole-cell patch-clamp recording from *tdTomato^PV^* cells. (**e**-**h** as in **a**-**d** respectively) for *tdTomato*^PV^ cells. Data are represented as mean ± SEM. One-way ANOVA or Two-way repeated measure ANOVA followed by Tukey test. * p < 0.05, ** p < 0.01.

Finally, we examined the involvement of Kv1.1 channels known to be involved in the delay of the first action potential onset and intrinsic excitability ^38^. To do so, we applied DTX-K, a selective blocker of Kv1.1, at 0.1 µM ^33^ after adding quinpirole, and measured DTX-K insensitive and DTX-K sensitive currents. In SST-INs, when evaluated at −40 mV, the effects of DTX-K are occluded by quinpirole while at −30 mV DTX-K is still able to decrease K^+^ conductance, suggesting a role for other members of the Kv1/2 families (**Supplemental Figure 3a**). In contrast, in the PV-INs, the quinpirole-induced increase in K^+^ current is blocked in the presence of DTX-K at both voltages indicating that D2R activation enables the modulation of Kv1.1 channel (**Supplemental Figure 3b**).

### D2R-mediated changes in SST-INs and PV-INs excitability rely on distinct D2R signaling

D2R signal through both G protein-dependent and non-canonical β-arrestin-dependent pathways. We therefore investigate whether quinpirole-induced modulation of SST-INs and PV-INs excitability relied on D2R/G-protein and/or D2R/β-arrestin signaling. Incubation of organotypic slices with pertussis toxin (PTX, 500 nM, ∼18h incubation) failed to prevent the increased intrinsic excitability, the decreased spike latency and ISI induced by quinpirole in SST-INs (**Figure 4a-d**). In contrast, GSK3 inhibition (SB-216763, 1 mM intrapipette) totally blunted D2R-mediated changes in SST-INs excitability (**Figure 4a-d**). Conversely, decreased PV-INs intrinsic excitability observed following bath application of quinpirole was not observed in presence of PTX while it remained unaffected by GSK3 inhibition (**Figure 4e-h**). Importantly, PTX and SB-216763 had no effect *per se* on spike number (**Figure 4b** and **e**). The observed effects were not the result from lack of efficacy of the drugs since SB-216763 reversed the increased excitability induced by quinpirole in mossy cells (Etter & Krezel, 2014) (**Supplemental Figure 4a, b**) and PTX prevented the effect of baclofen on CA3 pyramidal cells ^40^ (**Supplemental Figure 4c, d**). Together, these results indicate that D2R/β-arrestin signaling mediates the effect of D2R in SST-INs whereas in PV-INs D2R signals preferentially through the canonical pathway

**Figure 4:**
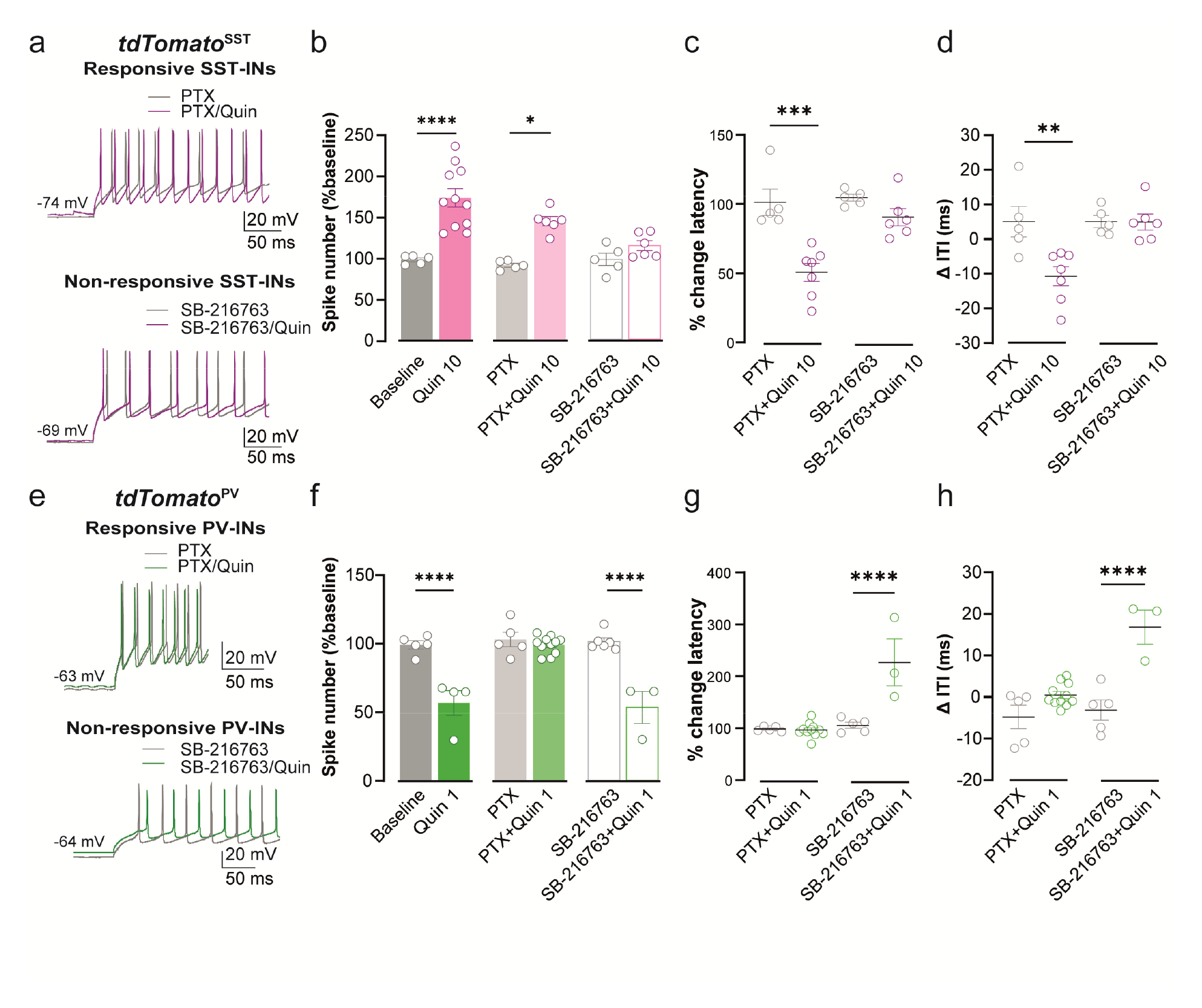
D2R activation signals through different intracellular pathways depending on the SST-INs or PV-INs. (**a**) Example traces of SST-INs showing the firing traces prior to and after quinpirole (10 µM) with PTX treatment (upper) or SB-216763 in the patch pipette (lower). (**b**) Mean responses corresponding to the last three min epoch of quinpirole 10 µM after different conditions. (**c**) Comparison of quinpirole-mediated latency change in presence of PTX or SB-216763. (**d**) Comparison of quinpirole-mediated variability of ITI in presence of PTX or SB-216763. (**e-h** as in **a-d** respectively) for PV-INs. Data are represented as mean ± SEM. One-way ANOVA followed by Tukey test. * p < 0.05, ** p < 0.01, *** p < 0.001, **** p < 0.0001.

### D2R activation modulate theta oscillations in the hippocampal CA1 area

To determine whether D2R signaling modulates hippocampal network, we then assessed the consequence of D2R activation on theta rhythm, a rhythmic activity of the network considered as the “on-line” state of the hippocampus and that can be artificially induced in organotypic slices ^41, 42^. We therefore investigated the consequence of D2R activation on methacholine-induced theta oscillation in CA1 pyramidal cells of organotypic hippocampal slices from C57Bl/6J mice (**Figure 5a**). Application of methacholine (MCh; 500 nM) during 20 min activates the hippocampal network (**Figure 5b-e**). Although the frequency of theta oscillations remained unchanged (**Figure 5c**), their occurrence was significantly increased in presence of quinpirole as revealed by the increased the number of theta oscillation (**Figure 5d**) and the mean time of each oscillation (**Figure 5e**). The effect of D2R on methacholine-induced theta oscillations were prevented in the presence of the D2R antagonist raclopride (40 µM) (**Figure 5b-e**). No theta oscillations were induced following the application of quinpirole in the absence of MCh, corroborating the role of dopamine as a neuromodulator (**Figure 5b-e**).

**Figure 5:**
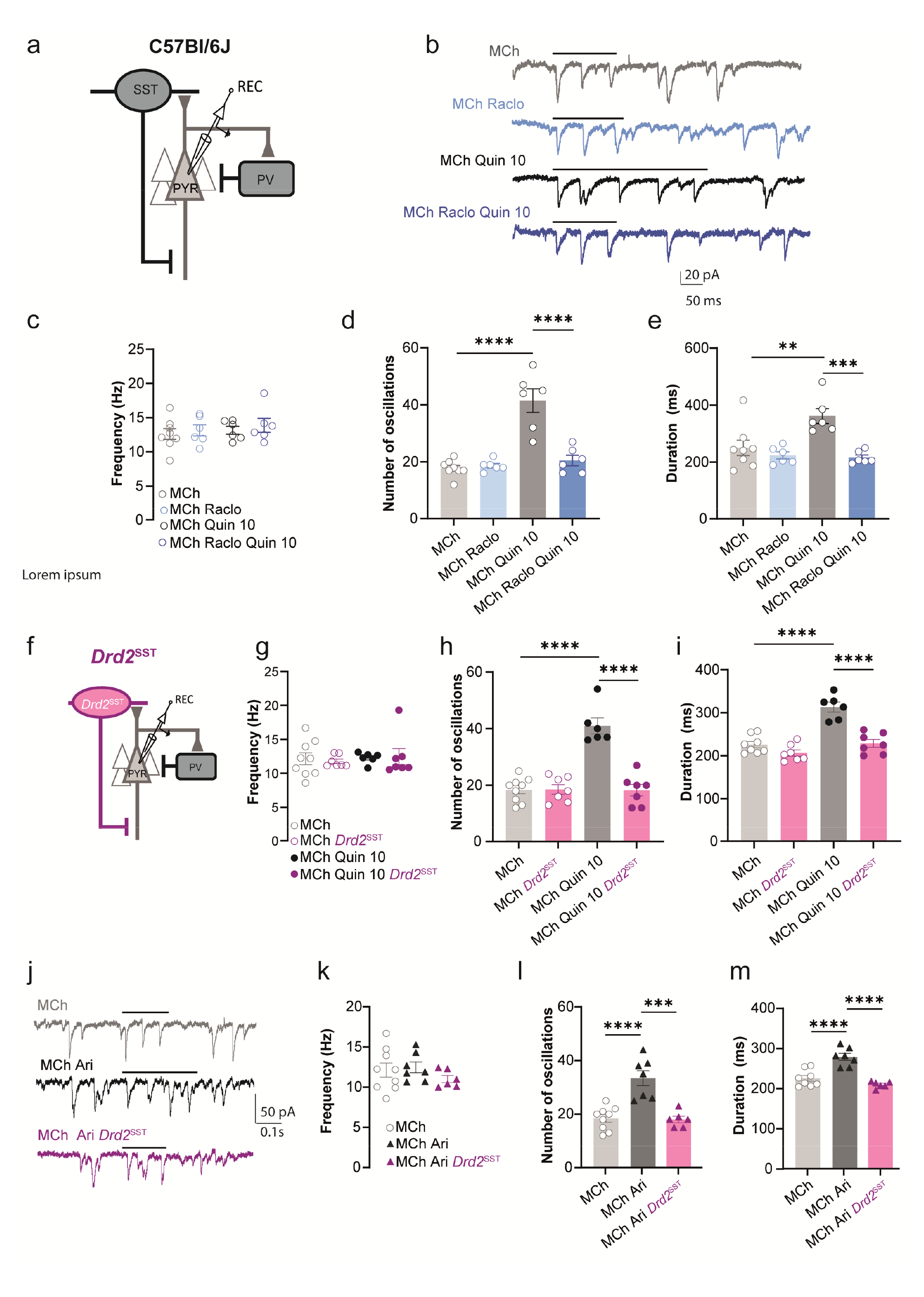
Modulation of theta oscillations in the hippocampal CA1 area by D2R activation is mediated by SST-INs. (**a**) Schematic representation of CA1 network in C57Bl/6J. (**b**) Representative traces of MCh-induced theta oscillation recorded in CA1 PC in different conditions. Black lines represent theta oscillation period. (**c**) Comparison of the frequency of each period of oscillation, (**d**) of the number of episode/min and (**e**) of the duration of each period of oscillation in MCh (Grey), MCh/raclopride (Blue), MCh/quinpirole (Dark Grey) and MCh/raclopride/quinpirole conditions (Dark blue). (**f**) Schematic representation of CA1 network in the *Drd2*^SST^mouse line. (**g**-**i** as in **c**-**e** respectively) from control and *Drd2*^SST^organotypic slices. (**j**) Example traces of MCh-induced theta oscillation prior to and after aripiprazole in control and *Drd2*^SST^ organotypic slices. Black lines represent period of theta oscillation. (**k**-**m** as in **g**-**i** respectively). Data are represented as mean ± SEM. One-way ANOVA followed by Tukey test. ** p < 0.01, *** p < 0.001, **** p < 0.0001.

Inhibitory network comprising PV-INs and SST-INs is important for the generation of these oscillations ^31, 43, 44^. We therefore investigated whether the modulatory role of D2R on CA1 theta oscillations is mediated by D2R signaling within SST-INs and/or PV-INs. For this purpose, we generated conditional transgenic mouse lines in which the gene encoding *Drd2* is invalidated in SST- (*Drd2*^SST^) and PV-INs (*Drd2*^PV^). D2R-mediated changes in SST-INs and PV-INs excitability were abolished in *Drd2*^SST^ and *Drd2*^PV^ mice confirming the lack of functional D2R in SST-INs and PV-INs (**Supplemental Figure 5**). Then, we investigated whether the lack of D2R in SST-INs and/or PV-INs impaired MCh-induced theta oscillations in CA1 pyramidal cells in organotypic hippocampal slice cultures from *Drd2*^SST^ and *Drd2*^PV^ mice (**Figure 5** and **Supplemental Figure 6**). No changes in the frequency, the number and duration of theta oscillations were observed in SST-INs and PV-INs lacking D2R (**Figure 5f-i** and **Supplemental Figure 6a-d**). In contrast, the ability of quinpirole to increase the number of oscillations and the duration of episodes was abolished in *Drd2*^SST^, but not in *Drd2*^PV^ mice (**Figure 5h, i** and **Supplemental Figure 6c, d**). Together, these results demonstrate that the effects of quinpirole on methacholine-induced rhythmic network activity require intact D2R signaling in hippocampal SST-INs.

### Aripiprazole modulates theta oscillations through SST-INs D2R signaling

Abnormal neural oscillations and synchrony constitute a hallmark of schizophrenia whose symptoms are primarily treated with atypical antipsychotic drugs, such as aripiprazole ^45^. In contrast to typical antipsychotics, aripiprazole presents a unique pharmacological profile by acting as a D2R partial agonist that displays biased signaling via β-arrestin ^45^. We therefore tested whether, similarly to quinpirole, bath application of aripiprazole (10 µM) could modulate MCh-induced theta oscillations. As shown in Figure 5, aripiprazole increased the number and duration of oscillations induced by MCh without altering their frequency (**Figure 5j-m**), these effects being not observed in *Drd2*^SST^ mice (**Figure 5j-k**). Altogether, these data indicate that modulation of theta oscillation by aripiprazole relies on intact D2R/ β-arrestin signaling in SST-INs.

## Discussion

Here, we found that D2R activation mediates opposite modulation of SST-INs and PV-INs excitability. These D2R-mediated changes rely on distinct intracellular pathways, involving the non-canonical β-arrestin-dependent pathway in SST-INs and the G protein-dependent pathway in PV-INs. Finally, our study also revealed that D2R activation modulates hippocampal theta oscillations through SST-INs D2R/β-arresting signaling.

The ability of D2R activation to modulate intrinsic excitability is known since decades. For instance, pharmacological D2R stimulation reduces the excitability of a wide range of neuronal population including principal neurons (both glutamatergic and GABAergic) as well as distinct classes of interneurons ^46–50^. Conversely, D2R activation increases PV-INs excitability in layer V of prefrontal and motor cortices ^51–53^, as well as in prefrontal cortex layer V pyramidal neurons, hippocampal glutamatergic hilar mossy cells and ventrobasal thalamic neurons ^39, 54, 55^. Our results indicating that D2R activation leads to opposite effects on hippocampal SST-INs and PV-INs excitability further illustrate that modulation of spike discharges frequency by D2R differs depending on brain areas studied, and cannot be predicted solely on the basis of the molecular identity of the recorded cells.

Control of neuronal excitability in response to D2R activation relies on coordinated modulation of distinct ion channels through multiple signaling mechanisms, which vary depending on the cell-types. Growing evidence suggest that the involvement of voltage-dependent K^+^ channels can finely regulate the neuronal excitability network ^33, 35^ which is frequently associated to D2R-mediated changes in excitability. Indeed, regulation of multiple K^+^ currents, including but not limited to inwardly rectifying K currents, "leak" currents and slowly inactivating K^+^ currents, contribute to reduced striatopallidal and dopamine neurons excitability ^47, 56, 57^. Similarly, regulation of K^+^ current participate to the increased spike discharge in Prefrontal cortex layer V pyramidal neurons and ventrobasal thalamic neurons ^54, 55^. Our results demonstrating that delayed rectifier K currents participate to D2R-modulated changes in hippocampal PV-INs and SST-INs excitability are in line with these previous studies. Interestingly, we found that D2R-mediated changes in PV-INs excitability rely on Kv1.1-containing K channels, while other members of Kv1 and/or Kv2 family channels are at play in SST-INs, the preferential enrichment of the *Kcna1* and the *Kcna2* transcripts in PV-INs and SST-INs respectively certainly accounting for such differences ^58^.

We then investigate whether the control of neuronal excitability relies on modulation of distinct Kv channels could be due to distinct signaling mechanisms depending on the cell-types. Interestingly, increasing evidence indicate that D2R mostly relies upon balanced G-protein and β-arrestin signaling ^59, 60^. We found that D2R-induced changes in PV-INs excitability strictly rely on the G protein-dependent pathway, while they required the non-canonical β-arrestin-dependent pathway in SST-INs, illustrating that multiple intracellular signaling mechanisms control D2R-mediated changes excitability. Interestingly, the transient decreased excitability observed in PV-INs suggests that the desensitization of D2R through its internalization could be an additional mechanism involved in the regulation of D2R-mediated changes in excitability. Whether other mechanisms are at play, depending on the localization of D2R in the somato-dendritic/ axon initial segments compartments ^61^ and/or the D2R isoforms (short vs long) expressed ^62–64^ will require future investigations.

Previous works support the role of D2R signaling in the regulation of hippocampal rhythms. Indeed, when administered systemically, the non-selective DA agonist, apomorphine shifts hippocampal theta waves to higher frequencies ^65^, and the D2R-like agonist, quinpirole, restores theta oscillations during periods of REM sleep in DA-depleted mice ^20^. Conversely, administration of the D2R antagonist, haloperidol, reduces hippocampal theta oscillations ^20, 65–67^. Here, we show that bath application of the D2R-like agonist, quinpirole, modulates methacholine-induced CA1 hippocampal theta oscillations. This effect is prevented by the selective D2R antagonist, raclopride, ruling out the potential involvement of D3R and/D4R. Importantly, our cell-type specific *Drd2* ablation allows us to identify that D2R-mediated enhancement of methacholine-induced rhythmic network activity relies on intact D2R signaling in hippocampal SST-INs but not PV-INs. These latter results further underscore the critical role of local SST-INs, as SST-INs strongly facilitate brain oscillatory activity, mainly in the theta range ^31, 43, 44, 68–70^.

Abnormal amplitude and synchrony of hippocampal oscillatory activity at theta frequency observed in schizophrenic patients has been proposed to contribute to cognitive impairments ^71–74^. Interestingly, we found that the atypical antipsychotic aripiprazole, also modulates hippocampal theta oscillations via SST-INs, thus identifying this class of interneurons as a key target through which DA exert its modulatory role in the hippocampus. Future studies will be needed whether disturbances of hippocampal oscillations observed in other neuropsychiatric disorders are also associated with altered functions of D2R in hippocampal SST-INs.

## Supplemental Figure Legends

**Supplemental Figure 1:**
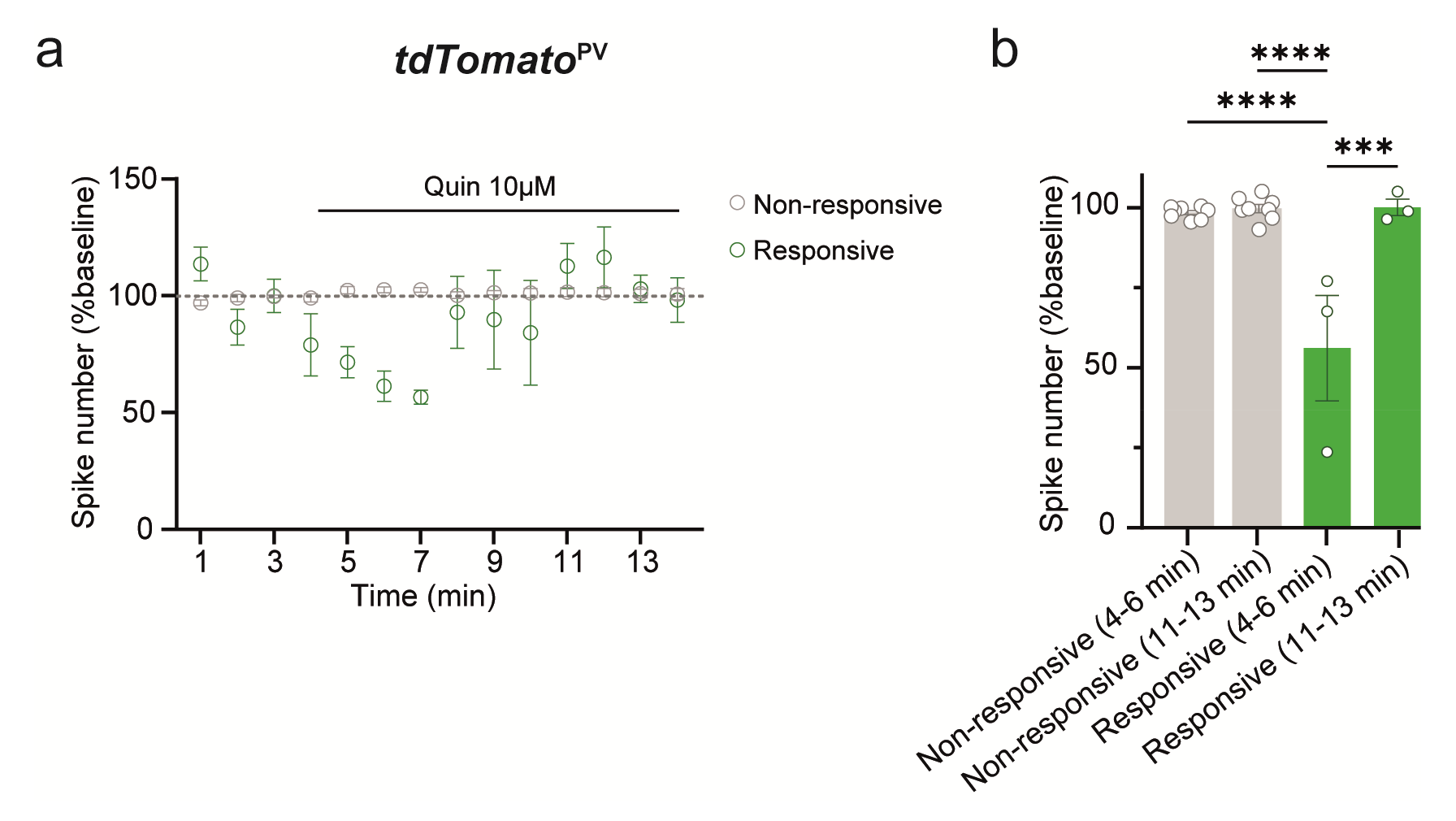
Application of quinpirole at 10µM induced an IE transient in PV-INs. (**a**) Time course of responses to depolarizing current steps prior and in response to quinpirole 10 µM of PV-INs. (**b**) Mean responses corresponding to the first and last three min of quinpirole application. Data are represented as mean ± SEM. One-way ANOVA followed by Tukey test. *** p < 0.001, **** p < 0.0001.

**Supplemental Figure 2:**
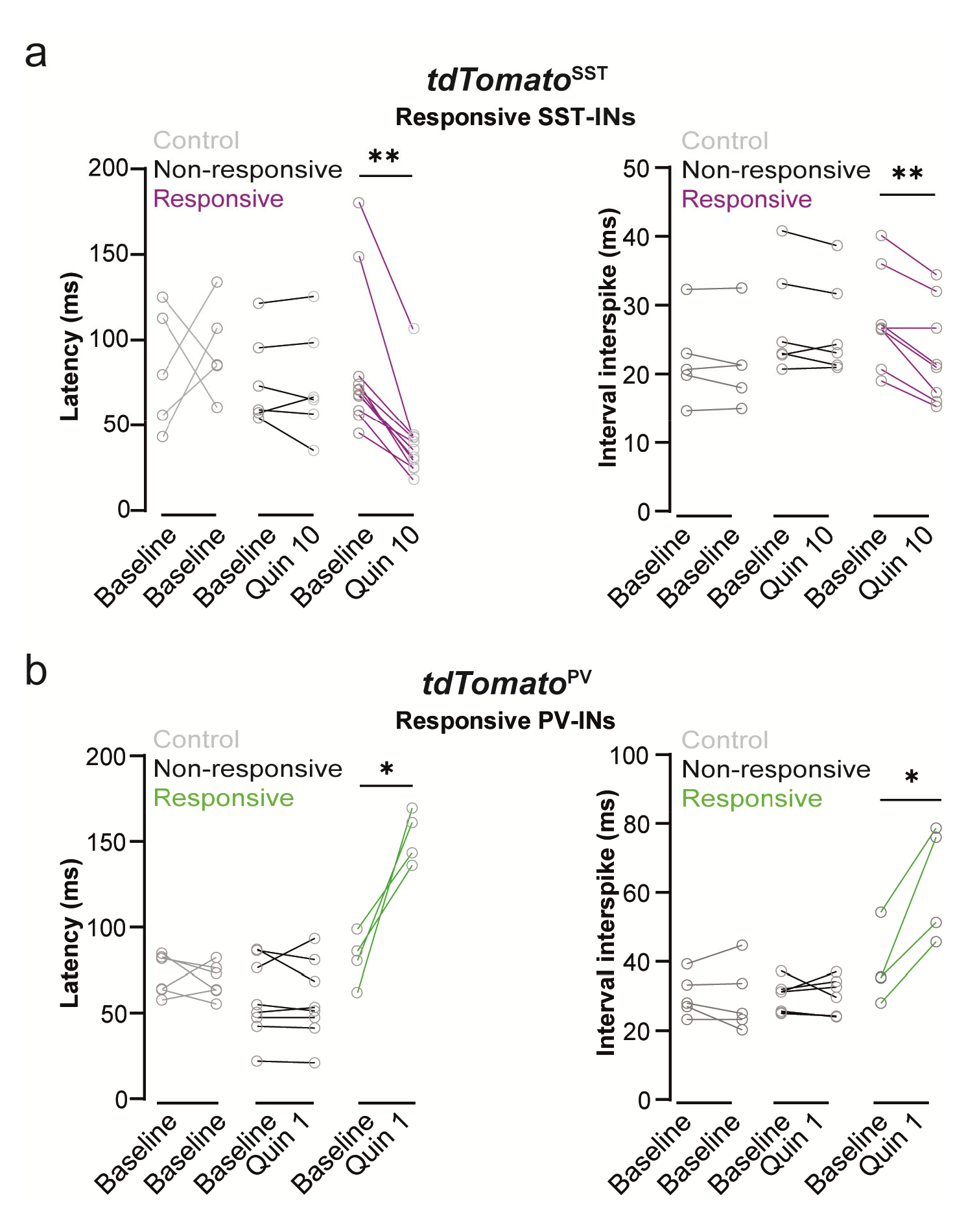
D2R activation alter spike latency and the interval interspike in SST-INs or PV-INs. (**a**) In *tdTomato*^SST^ neurons. Left, comparisons changes in latency between baseline/baseline or baseline/quinpirole. Right, summary of ITI (ms) before and after the application of quinpirole 10 µM. (**b** as in **a**) in *tdTomato*^PV^ cells. Unpaired t-test. * p < 0.05, ** p < 0.01.

**Supplemental Figure 3:**
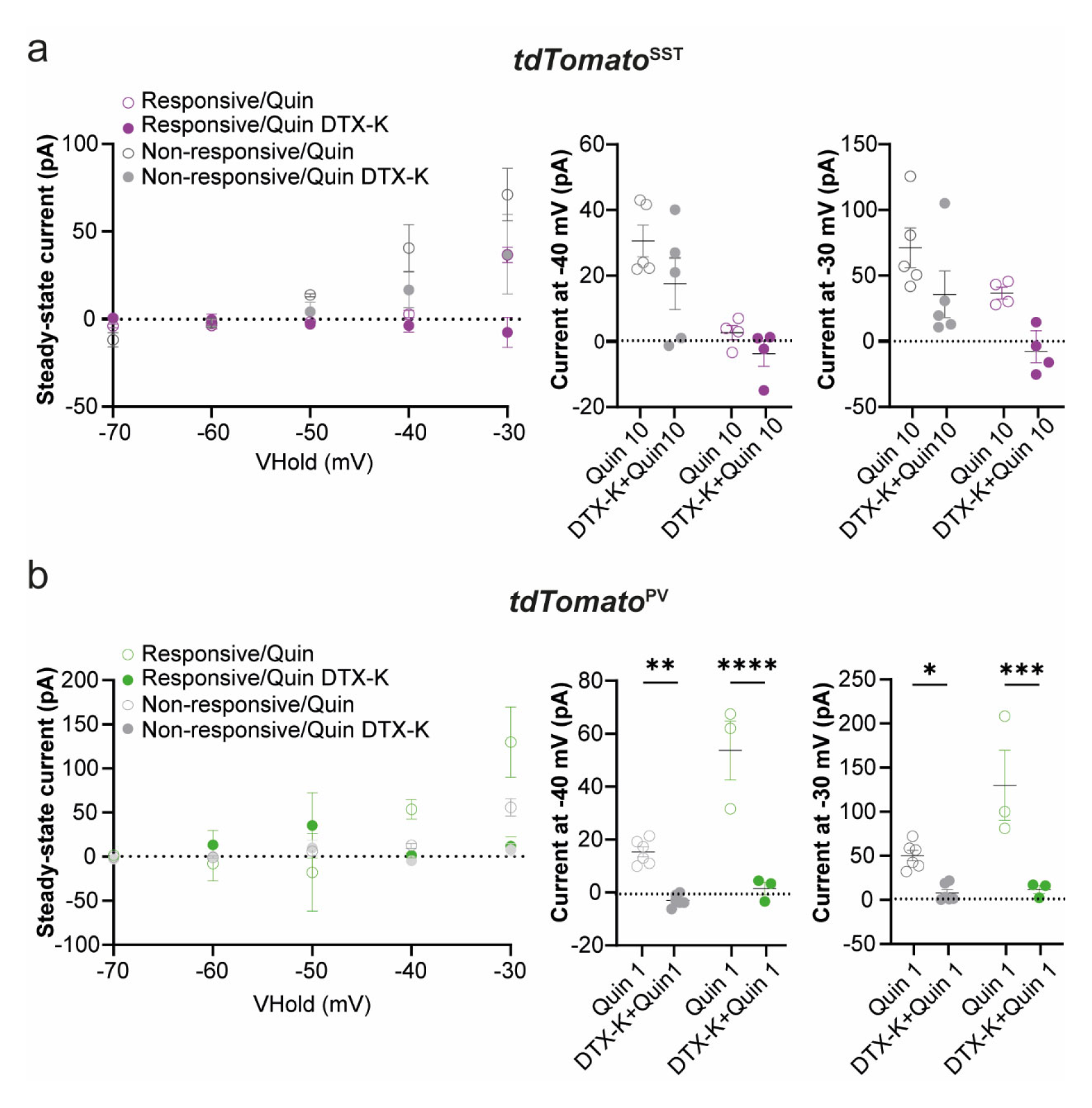
Implication of Kv1.1 channel in D2R activation-mediated changes in active membrane properties in SST-INs and PV-INs. **(a)** Left, I-V curve of leak-subtracted current for *tdTomato*^SST^ cells from −70 to −30 mV after application of quinpirole or quinpirole/DTX-K. Mean of leak-subtracted steady state at −40 mV in SST-INs after application of quinpirole 10 µM with or without DTX-or at −30 mV. Left part: Ratio of decrease for SST-INs at −30mV after DTX-K application. (**b** as in **a**) for *tdTomato*^PV^ neurons after quinpirole 1 µM application with or without DTX-K at −30 mV. Data are represented as mean ± SEM. Unpaired t-test and Two-way repeated measure ANOVA followed by Tukey test. * p < 0.05, ** p < 0.01, *** p < 0.001, **** p < 0.0001.

**Supplemental Figure 4:**
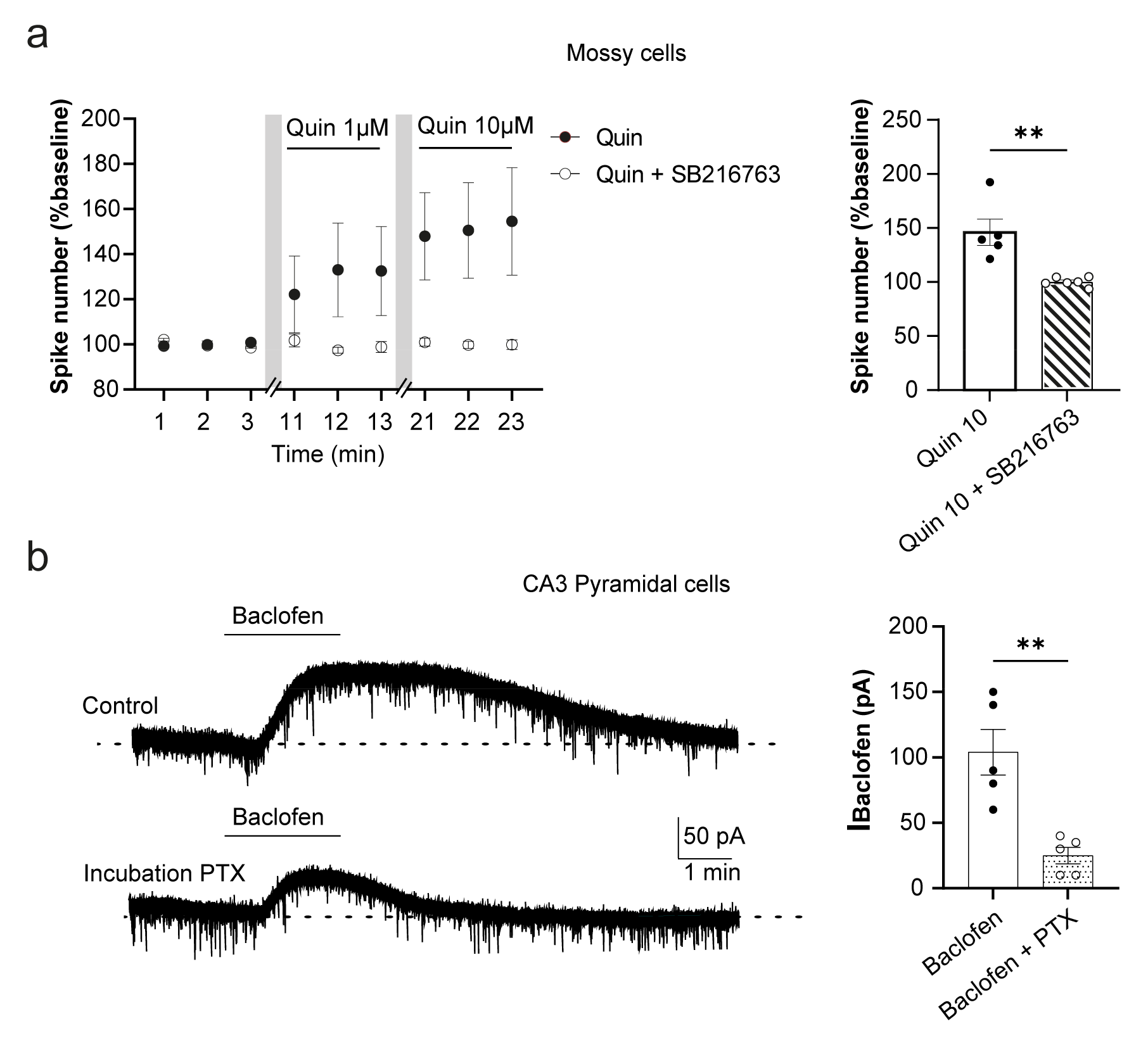
Excitability changes in mossy cells by Quinpirole application is reversed by SB-216763 and Baclofen induced-current are abolished by PTX treatment. **(a)** Left, time course of responses to depolarizing current steps prior and in response to quinpirole 10 µM with or without SB-216763 in mossy cells. Right, mean responses corresponding to the last three min epoch. (**b**) Representative voltage clamp trace of Baclofen inducing K^+^ current with or without PTX treatment in CA3 PC. Right, K^+^ current induced by baclofen with or without PTX treatment. Data are represented as mean ± SEM. One-way ANOVA followed by Tukey test. ** p < 0.01.

**Supplemental Figure 5:**
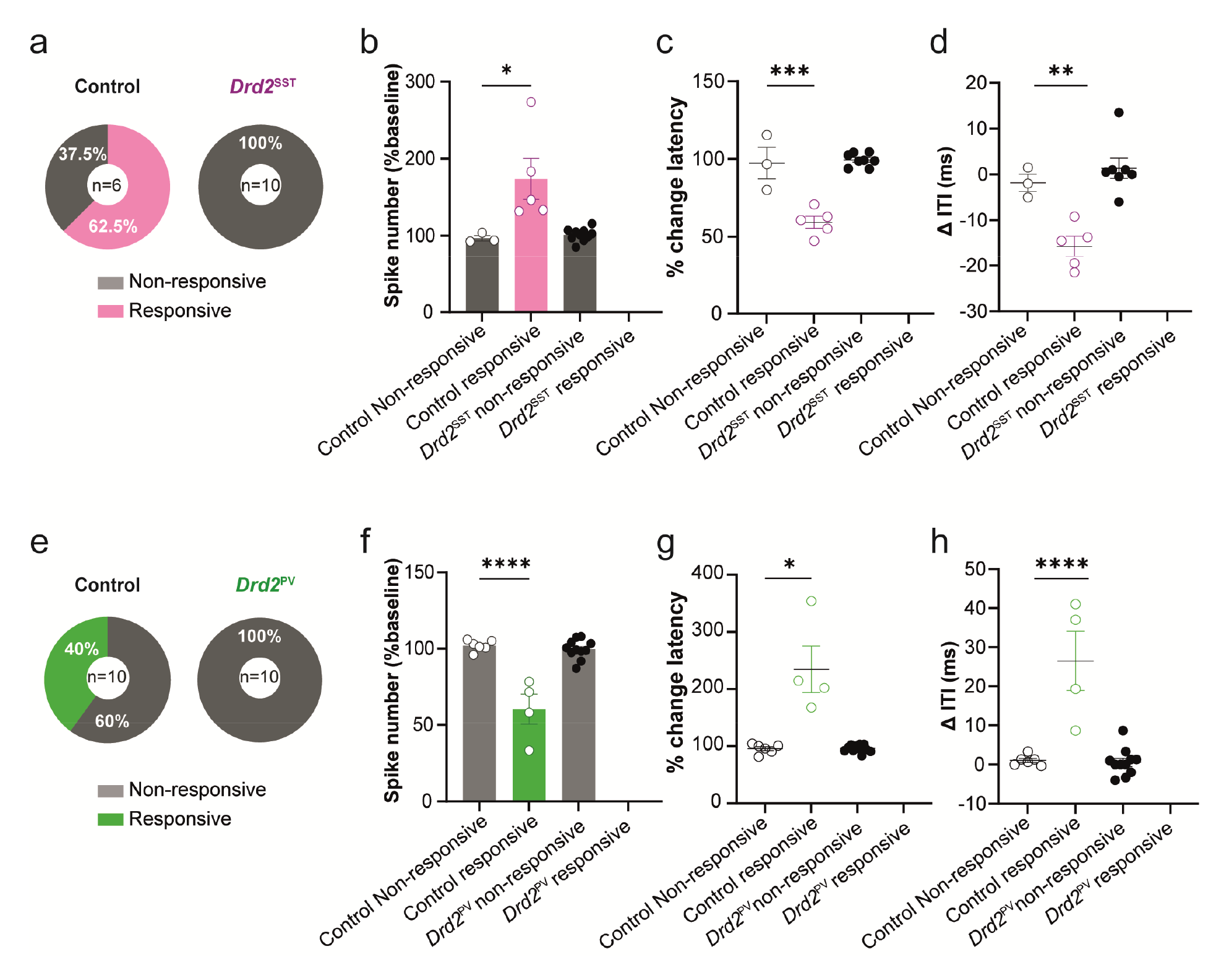
D2R-mediated changes in excitability are abolished in *Drd2*^SST^and *Drd2*^PV^. **(a)** Pie chart of SST-INs in response to quinpirole 10 µM in control littermate and *Drd2*^SST^ INs. **(b)** Mean responses corresponding to the last 3 min epoch of quinpirole 10 µM. (**c**) Comparison of normalization in latency to spike and (**d**) of variability of ITI from control and *Drd2*^SST^organotypic slices. (**e** as in **a**) of PV-INs in response to quinpirole 1µM in control and *Drd2*^PV^ organotypic slices. (**f**-**h** as in **b-d** respectively) for *Drd2*^PV^ organotypic slices. Data are represented as mean ± SEM. One-way ANOVA followed by Tukey test. * p < 0.05, ** p < 0.01, *** p < 0.001, **** p < 0.0001.

**Supplemental Figure 6:**
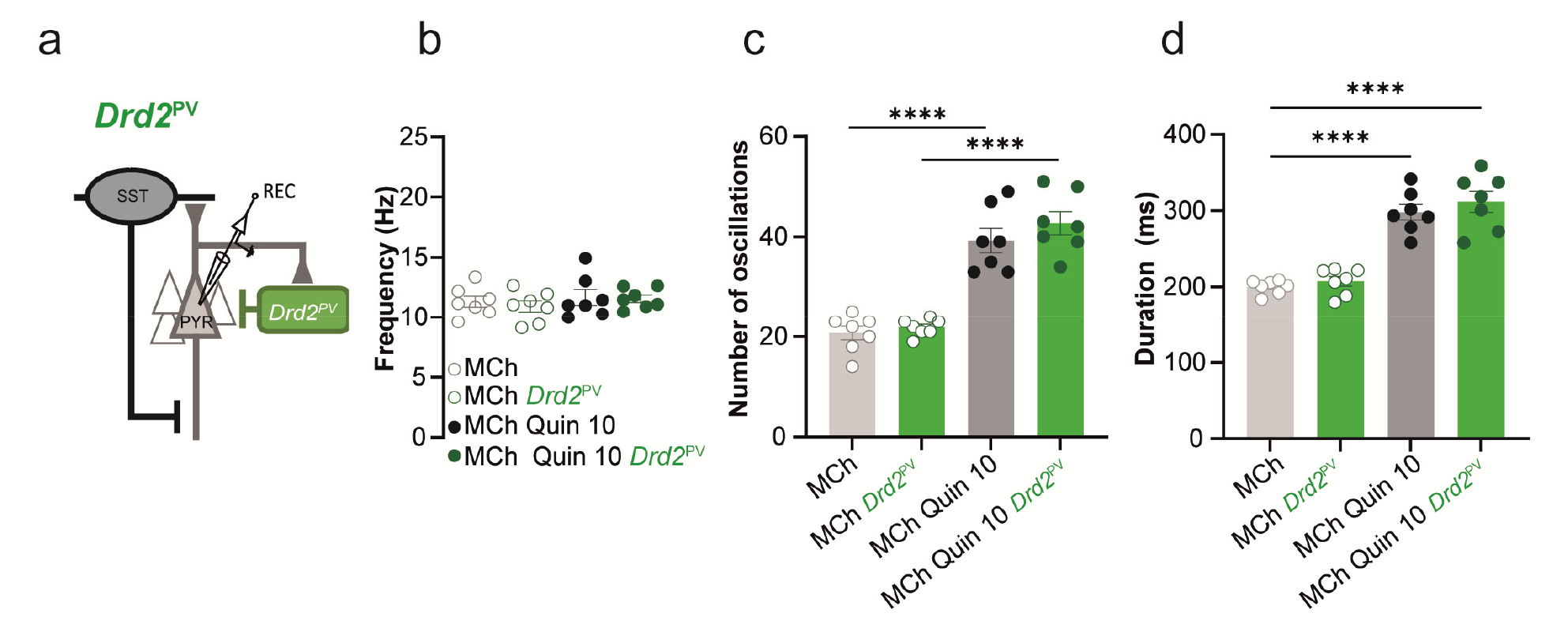
The increase of theta oscillation induced by D2R is not mediated by PV-INs. (**a**) Schematic representation of CA1 network in the *Drd2*^PV^mouse line. (**b**) Frequency of each period of oscillation in control and *Drd2*^PV^organotypic slices. (**c**) Number of episodes/min in control and *Drd2*^PV^ organotypic slices. (**d**) Duration of each period of oscillation in control and *Drd2*^PV^ organotypic slices. Data are represented as mean ± SEM. One-way ANOVA followed by Tukey test. **** p < 0.0001.

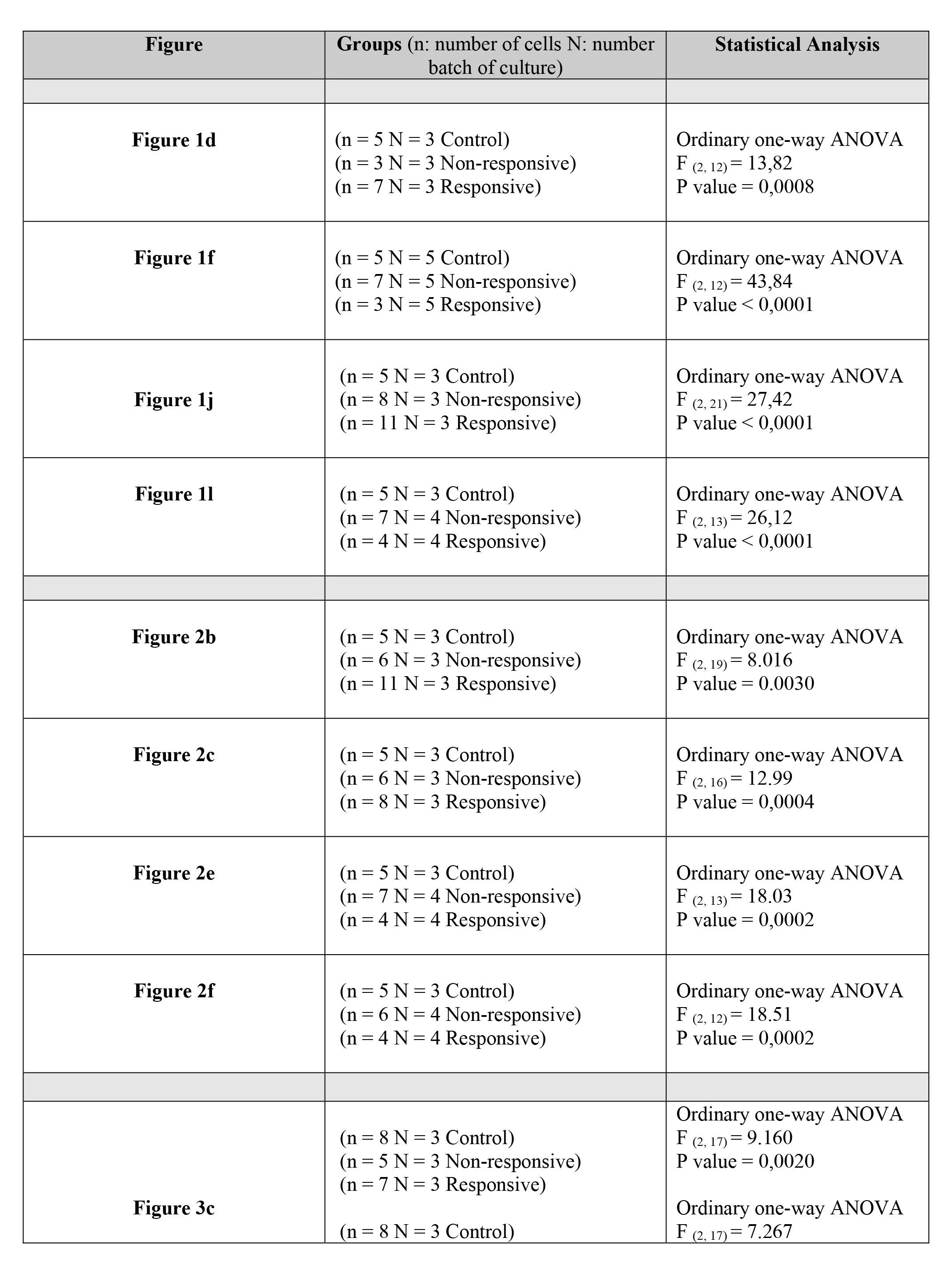
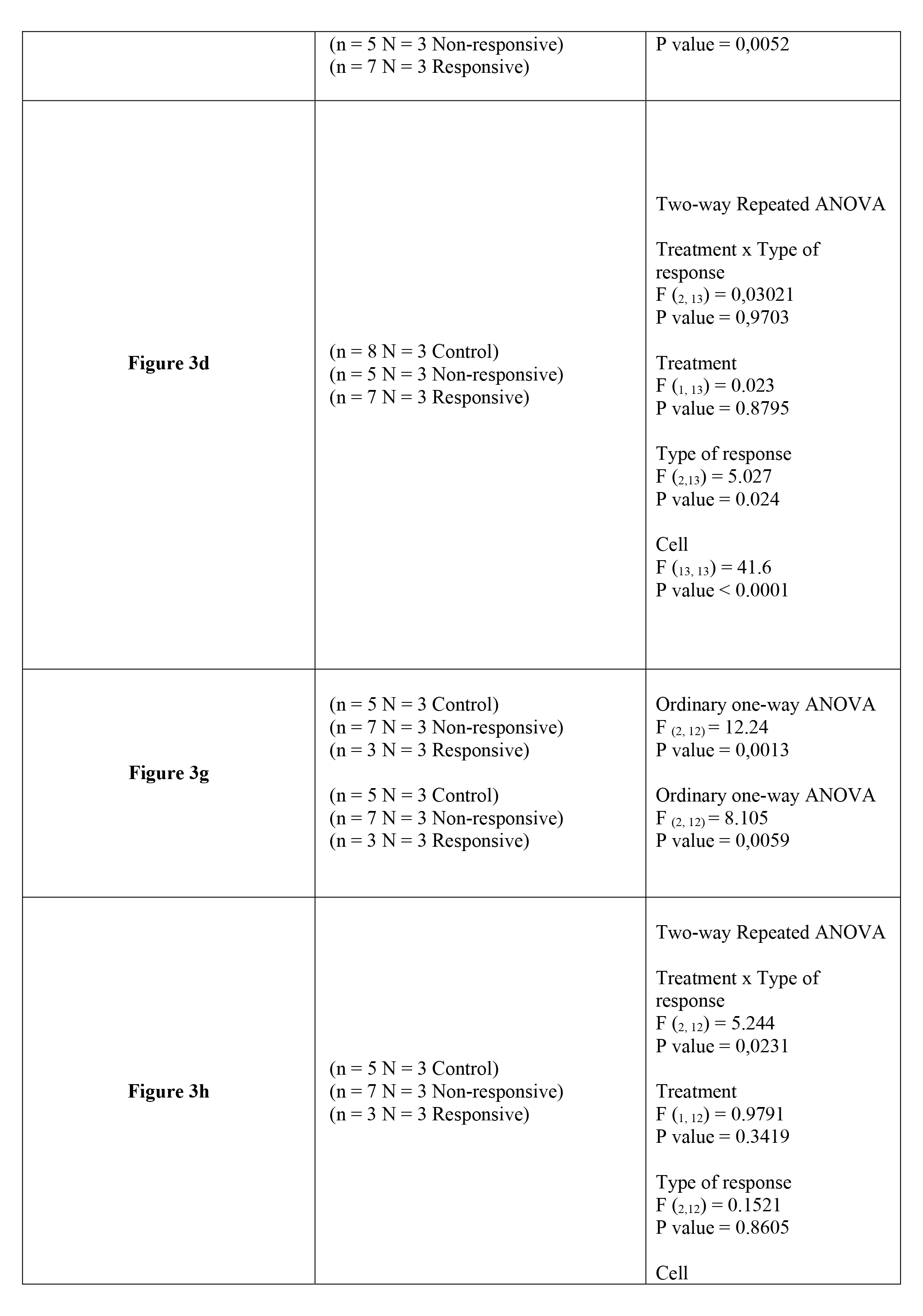
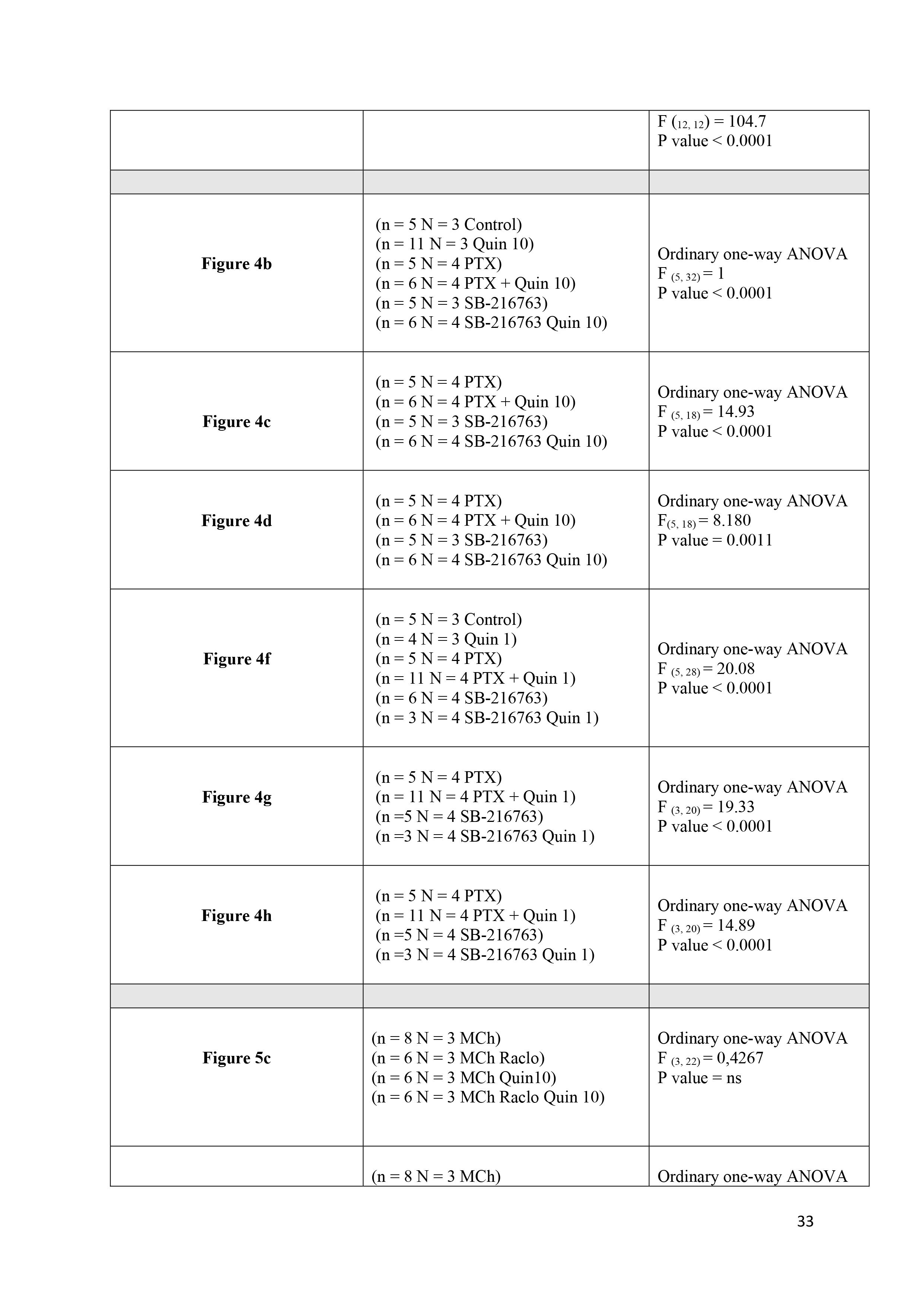
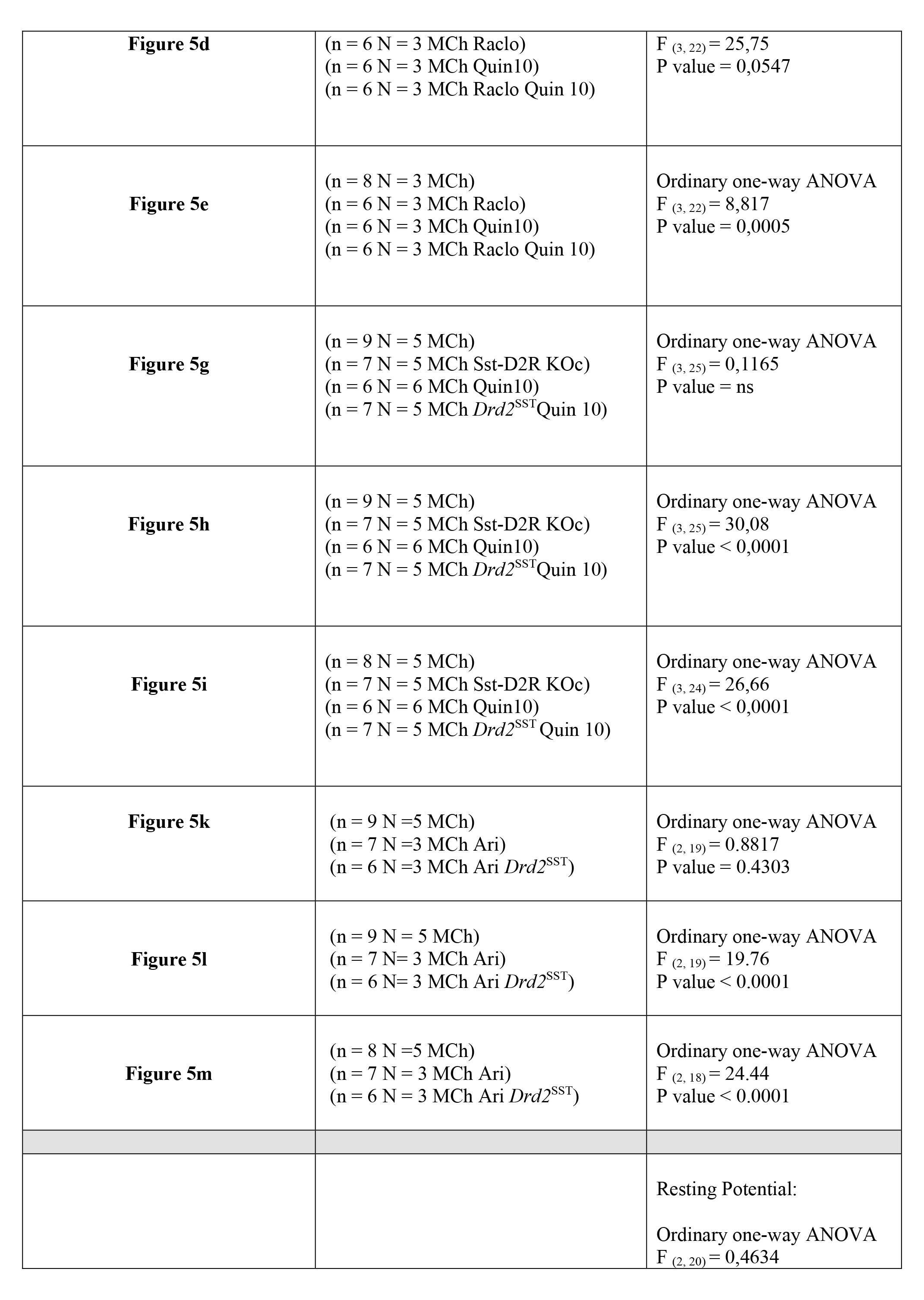
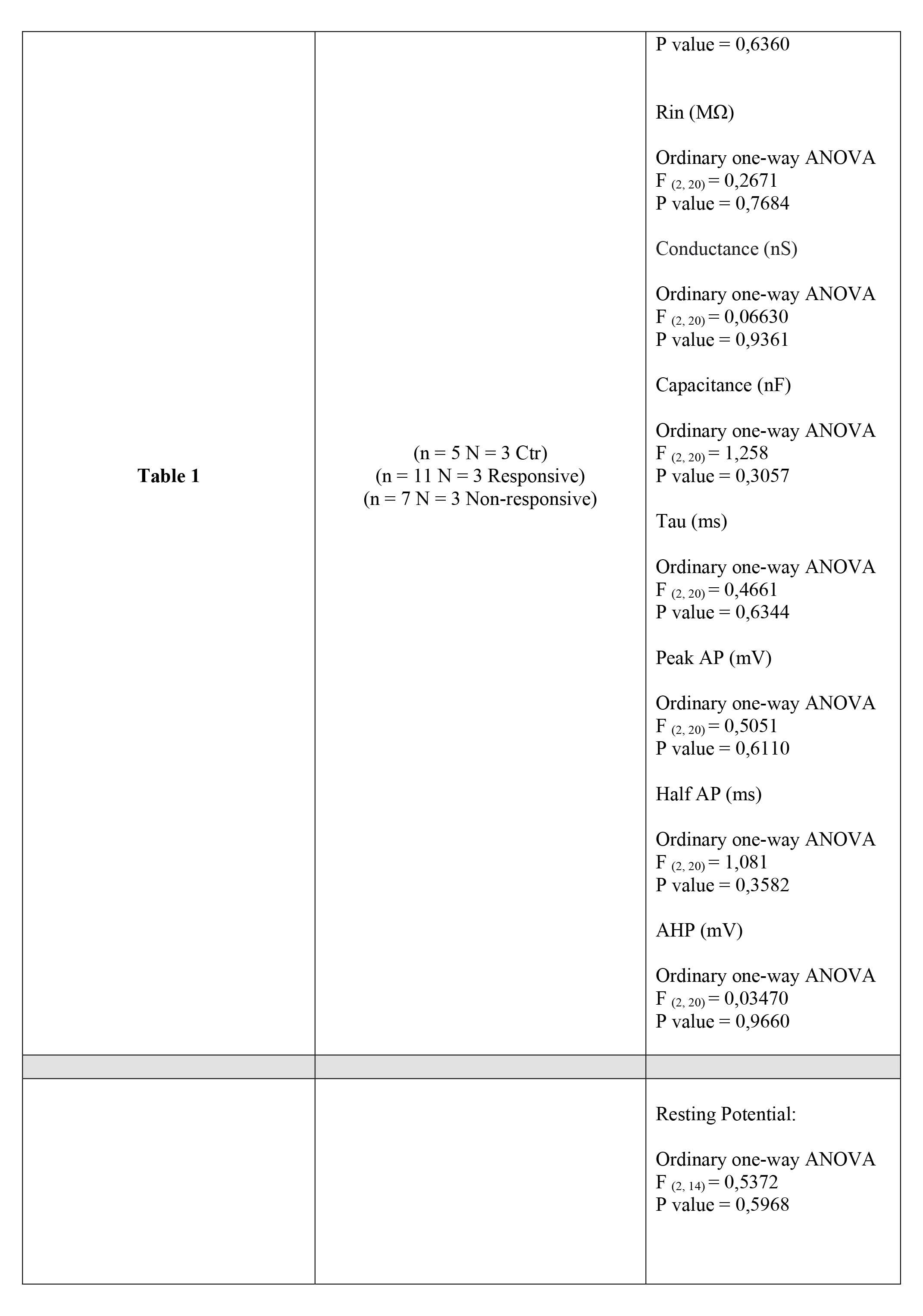
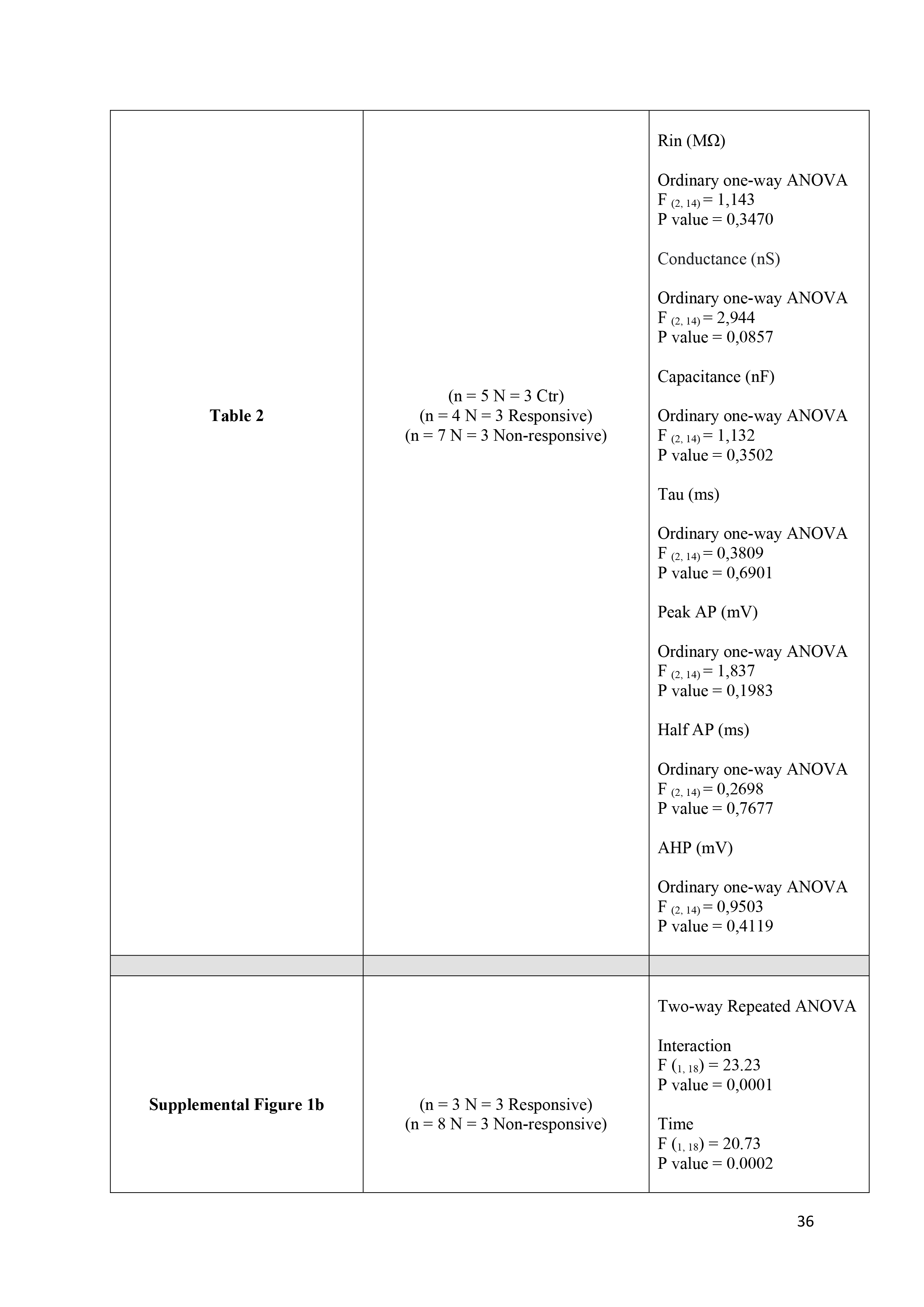
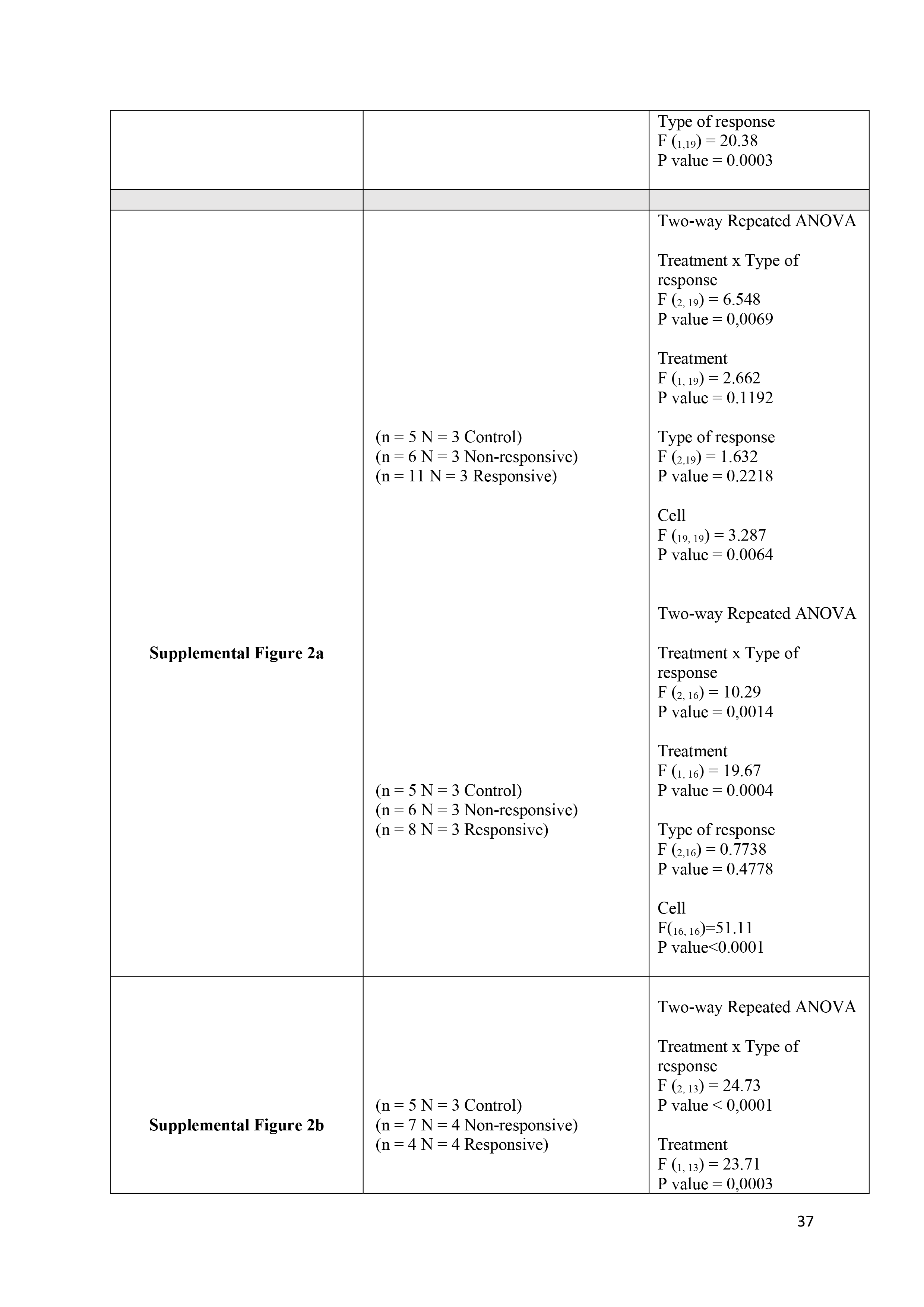
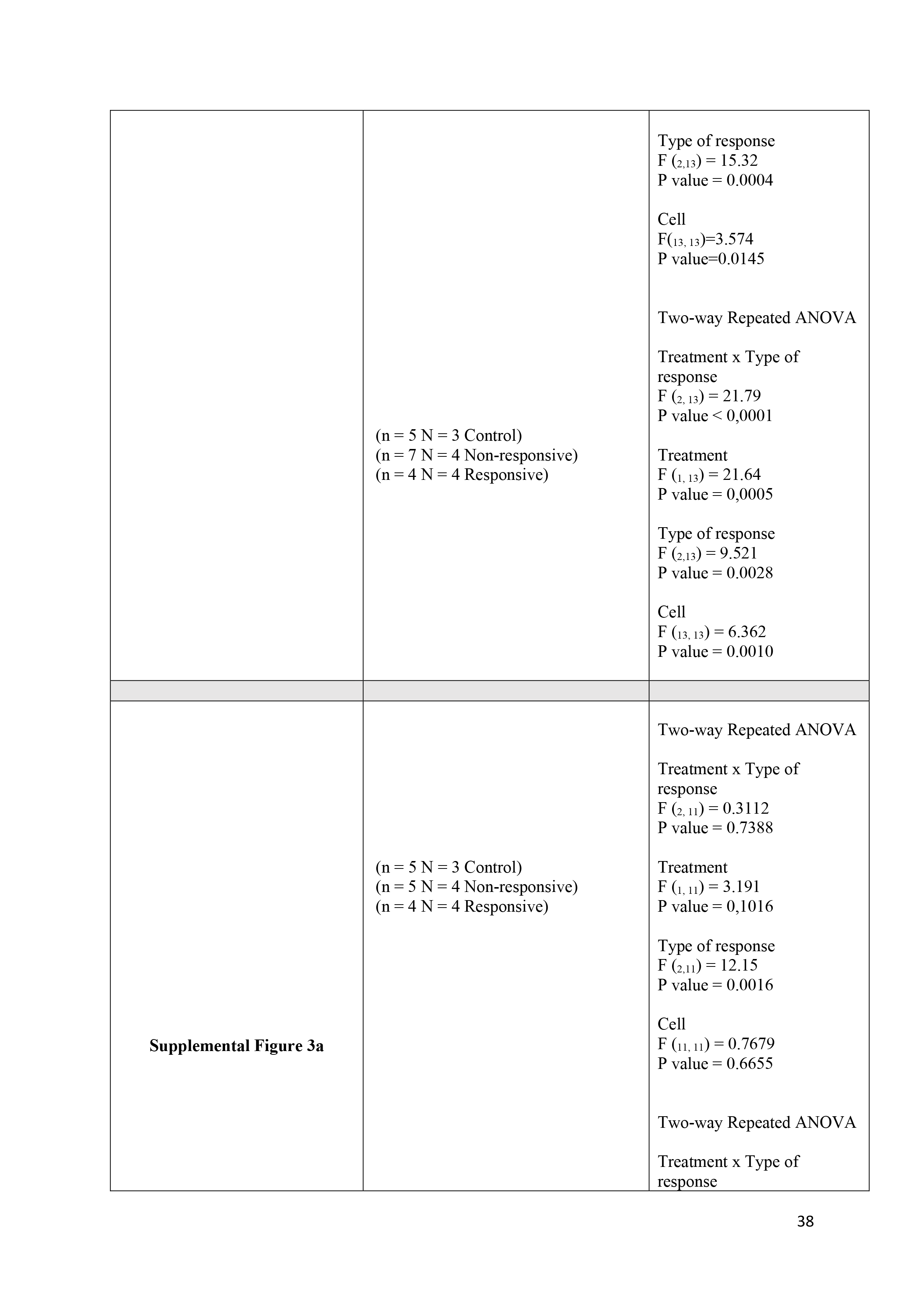
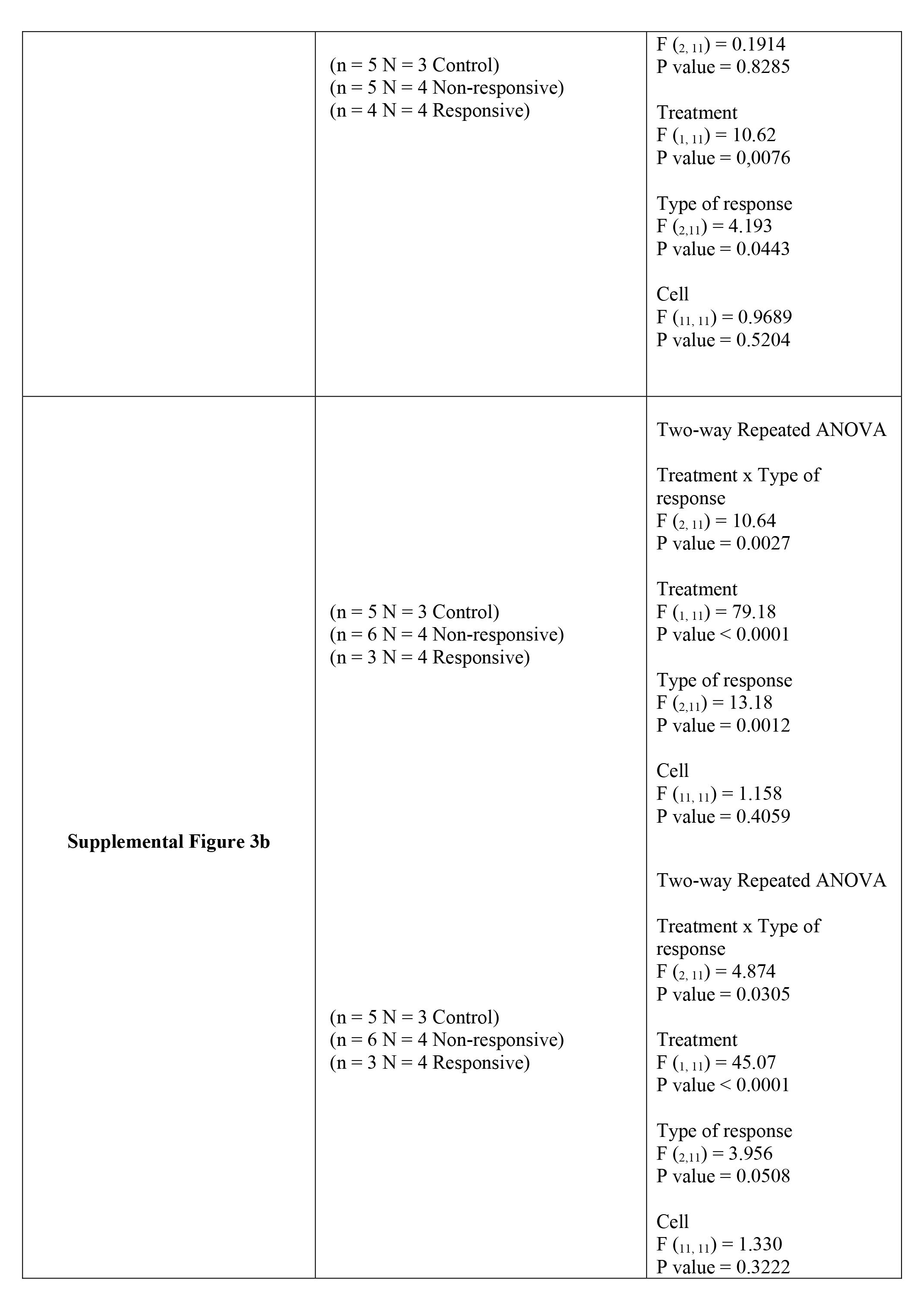
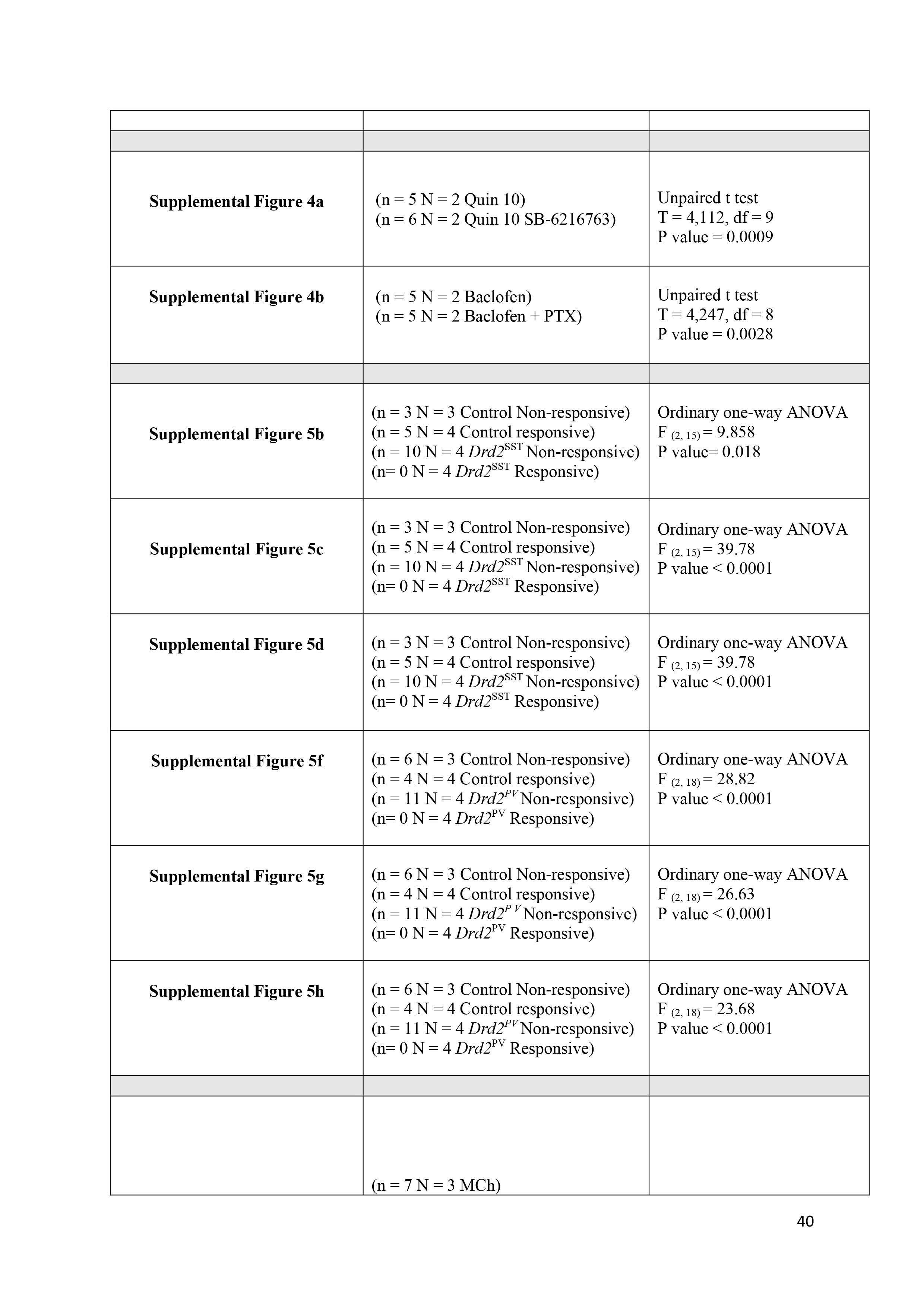
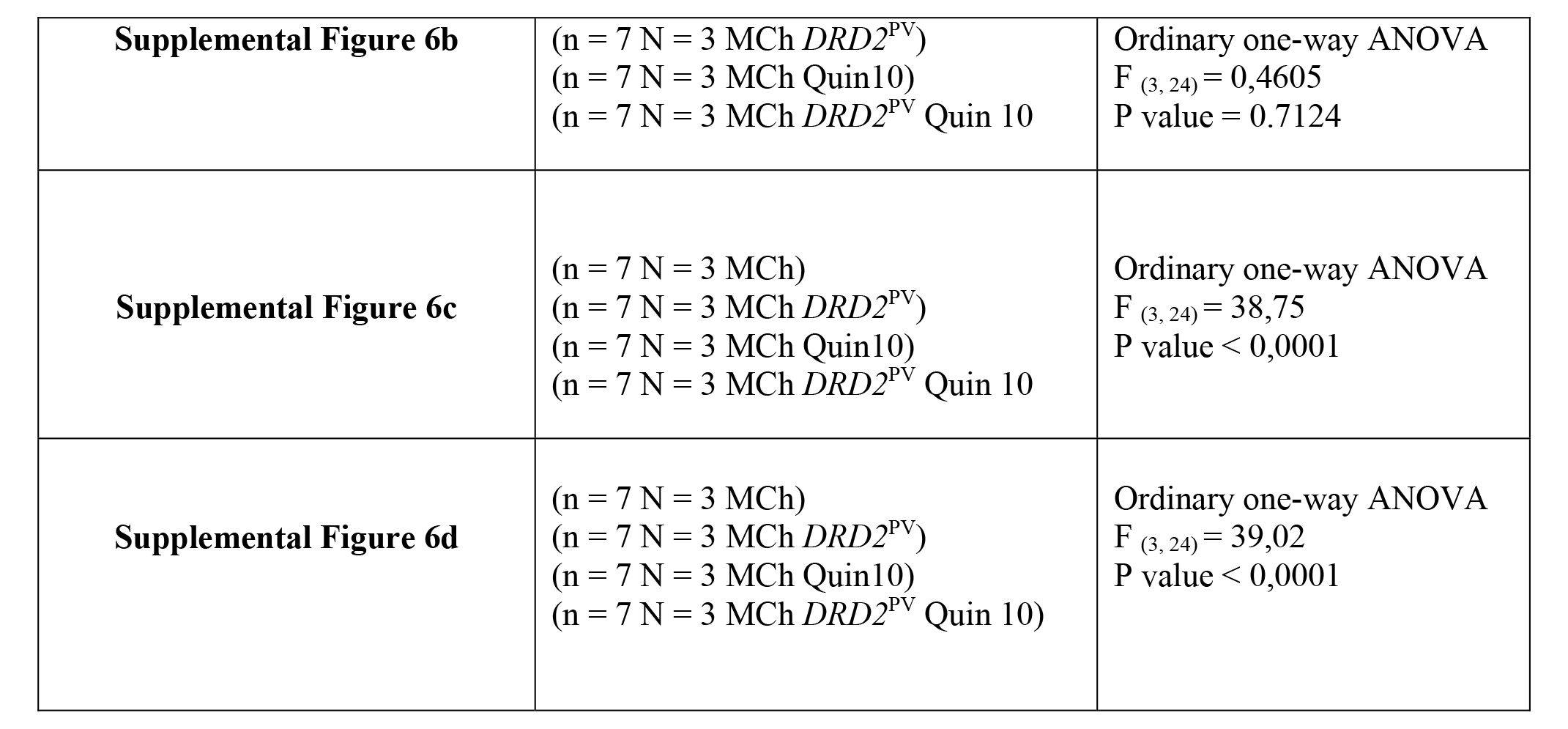
Supplemental Table.

